# WNT/β-catenin dependant alteration of cortical neurogenesis in a human stem cell model of SETBP1 disorder

**DOI:** 10.1101/2021.10.12.464034

**Authors:** Lucia F. Cardo, Meng Li

## Abstract

Disruptions of *SETBP1* (SET binding protein 1) on 18q12.3 by heterozygous gene deletion or loss-of-function variants cause SETBP1 disorder. Clinical features are frequently associated with moderate to severe intellectual disability, autistic traits and speech and motor delays. Despite *SETBP1* association with neurodevelopmental disorders, little is known about its role in brain development. Using CRISPR/CAS9 genome editing technology, we generated a SETBP1 deletion model in human embryonic stem cells (hESCs), and examined the effects of SETBP1-deficiency in in vitro derived neural progenitors (NPCs) and neurons using a battery of cellular assays, genome wide transcriptomic profiling and drug-based phenotypic rescue.

SETBP1-deficient NPCs exhibit protracted proliferation and distorted layer-specific neuronal differentiation with overall decrease in neurogenesis. Genome wide transcriptome profiling and protein biochemical analysis showed that SETBP1 deletion led to enhanced activation of WNT/β-catenin signaling. Crucially, treatment of the SETBP1-deficient NPCs with a small molecule WNT inhibitor XAV939 restored hyper canonical β-catenin activity and rescued cortical neuronal differentiation.

Our study establishes a novel regulatory link between SETBP1 and WNT/β-catenin signaling during human cortical neurogenesis and provides mechanistic insights into structural abnormalities and potential therapeutic avenues for SETBP1 disorder.

## INTRODUCTION

The cerebral cortex is the center of higher mental functions for humans and contains around 100 billion cells that account for about 76% of the brain’s volume. Normal cortical development involves a set of highly complex and organized events, including neural stem cell proliferation, neuronal differentiation and appropriate positioning and interconnection of both excitatory and inhibitory neurons(*1-3*). Abnormalities of cell proliferation or neurogenesis may cause malformations of the brain such as microcephaly, macrocephaly or cortical dysplasia. Cortical malformations and aberrant neural circuitry have been implicated as an important cause of neurological disorders such as intellectual disability, autism and developmental delay(*4-7*).

*SETBP1* gene is located at 18q12.3 and is associated with several neurodevelopmental disorders. SETBP1 haploinsufficiency due to heterozygous gene deletion or loss-of-function mutation cause SETBP1 disorder, a rare disorder with clinical features including expressive language impairment, intellectual disability, autistic-like traits, autism spectrum disorder (ASD), attention deficit hyperactivity disorder (ADHD), seizures, delayed motor skills and minor dysmorphic features amongst others(*8-14*). The disorder is also known as SETBP1 haploinsufficiency disorder or Mental Retardation Dominant 29 (MIM #616078). Its strong association with a phenotype of developmental delay with language disorder, makes SETBP1 a new candidate gene for speech disorders(*8, 12, 15, 16*). In contrast, point mutations of *SETBP1* result in SETBP1 gain-of-function due to impairment in its degradation(*17*) and causes a different disorder called Schinzel-Giedion syndrome (SGS). SGS is a severe multi-organ disorder characterized by distinctive facial features, profound neurodevelopmental and structural anomalies and higher prevalence of myeloid leukaemia(*18-20*). Despite its clear association with several neurodevelopmental disorders, the function of SETBP1 in the developing brain remains unknown.

Human embryonic stem cells (hESCs) offer an infinite cell source for the generation of neural progenitors (NPCs) and neurons, and have proved to be an invaluable *in vitro* model for studying human neurodevelopment and associated neurological disorders. HESCs provide an isogenic model with defined genetic background in which disease-associated mutations can be generated using genome editing tools, such as the state-of-the-art CRISPR/CAS9 genome editing technology(*21-23*). In this study, we generated a SETBP1 loss-of-function hESC model via CRISPR/CAS9 assisted gene targeting and investigated its impact on cortical neuronal differentiation.

## RESULTS

### Generation of SETBP1-deficient hESC lines

CRISPR/CAS9 genome editing technology was employed to generate a hESC model of *SETBP1* deletion. A classic gene targeting vector was designed for creating a defined deletion by homologous recombination in exon 4, which encodes two AT hooks, SKI homologous region and SET binding domain of SETBP1 protein. To guide Cas9 cleavage of the target DNA, three gRNAs were co-transfected with the donor template. Independent hESC clones were screened for homologous recombination by PCR followed by Sanger sequencing (Fig. 1A, 1B and Fig. S1A). Three heterozygous (SETBP1+/-) lines were obtained from the first round of gene targeting, which introduced an early stop codon in the mutant allele and is predicted to produce a truncated protein of 475 of the 1596 amino acid full protein sequence (Fig. 1C). One of this lines (HET1) was subjected to a second round of editing using the same gRNAs without the donor template, yielding several independent clones containing a 5bp deletion in the other allele (ie. Homozygous SETBP1 mutant lines, Fig. S1B and Fig. S1C). This 5bp deletion introduced a premature stop codon in the second allele and is predicted to produce a truncated protein of 1220 amino acid (www.expasy.org. Fig. 1C).

**Fig. 1.**
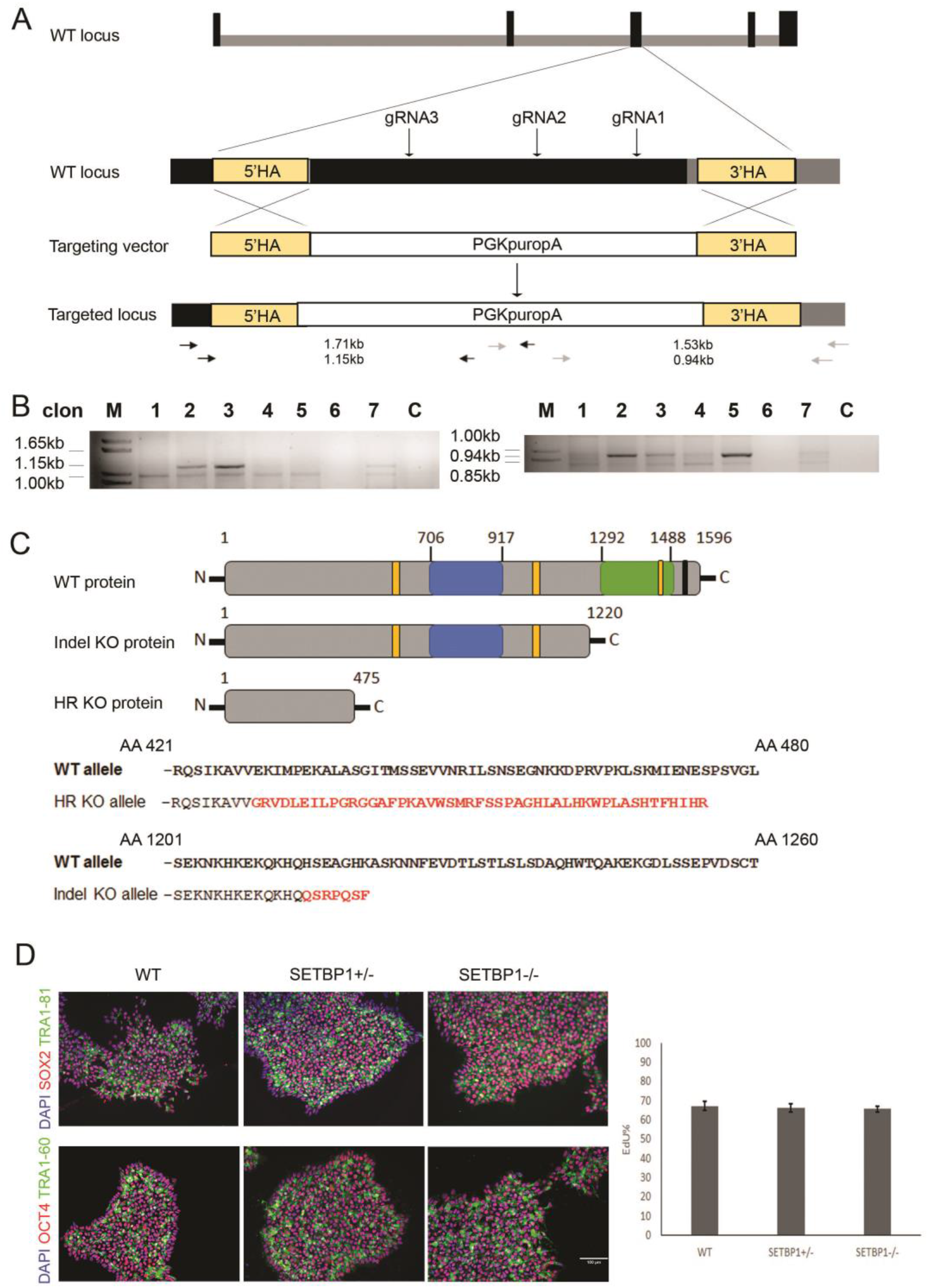
Generation of the SETBP1-deficient hESC lines. (**A**) Schematic illustration of the wild type (WT) *SETBP1* locus and targeting strategy. Exons are shown in black and introns in grey. The three gRNAs targeting exon 4 are indicated in arrows. The homologous arms (HA) corresponding to exon 4 and part of intron 4/5 are indicated in yellow, which are flanked by a PGKpuropA selection cassette in the targeting vector. The positions of the two nested PCR primer pairs for screening homologous recombination (HR) at the 5’ and 3’ are indicated in black and grey arrows, respectively. (**B**) Agarose gels showing the PCR amplicon from the targeted clones (lanes 2, 3, 7). (**C**) SETBP1 protein aligned with predicted mutant SETBP1 proteins for HR -/- (475AA) and indel -/- (1220AA) alleles, respectively. yellow=AT hook domains, blue=SKI homologous region, green=SET binding domain, black=repeat domain. Amino acid sequence alignment of WT SETBP1 protein vs HR -/- allele and indel -/- allele are shown. Amino acids in red indicate the sequence different from the WT prior to the stop codon. (**D**) Representative immunostaining of WT, *SETBP1*+/- and *SETBP1*-/- clones for pluripotency markers SOX2 (red), TRA1-81 (green) and OCT3/4 (red) with DAPI counterstain. Bar graph shows the proportion of EdU^+^ in WT, SETBP1+/- and SETBP1-/- hESCs (P>0.05). Data presented as mean ± s.e.m of two independent experiments. AA: amino acid; N: NH2 terminal; C:COOH terminal. Scale bar: 50uM.

Three homozygous SETBP1 mutant hESC lines (KO1, KO2 and KO3, referred together as SETBP1-/-), along with two SETBP1+/-lines, were chosen for subsequent studies. The SETBP1 edited lines exhibited characteristic pluripotent stem cell (PSC) morphology, expressed pluripotency markers OCT4 and SOX2, and grew at a similar rate to that of H7 (Fig. 1D). Moreover, they have normal karyotype (46, XX) in 73-82% of total metaphases analysed.

### Loss of SETBP1 affects neural rosette size without compromising neural induction

SETBP1 is expressed in the ventricular zone of the developing mouse telencephalon (http://www.eurexpress.org) and highly expressed in human neocortex (http://hbatlas.org). Cortical differentiation was therefore chosen for investigating SETBP1 function. The SETBP1 deficient and isogenic control hESCs were induced to differentiate towards forebrain fate using a modified dual SMAD inhibition protocol as described previously (Fig. 2A)(*24, 25*). Efficiency of cortical fate commitment was analysed by antibody staining for pan neural stem cell markers (SOX2, NESTIN) and the forebrain and dorsal NPC markers (FOXG1, PAX6 and OTX2) at day 18-20 (Fig. S2A-B). The vast majority of cells, stained positive for PAX6, OTX2, SOX2 and NESTIN and the numbers of positive cells were comparable between the three genotypes, with the exception of FOXG1 that exhibited a 50% reduction in the SETBP1-/- cultures in comparison with WT levels. Marker expression at the protein level was supported by RT-PCR analysis, which revealed a rapid exit of the pluripotent state as demonstrated by downregulation of *OCT4* and *NANOG* and concurrent induction of *SOX1, SOX2, NESTIN, PAX6, OTX2*, and additional dorsal NPC gene markers *EMX2* and *GLI3* in a similar temporal pattern between genotypes. Consistent with immunostaining, we detected a decreased level of *FOXG1* and *SIX3* in SETBP1-/- cultures compared to the WT (Fig. S3A). Non-cortical transcripts such as *PAX7* (dorsal midbrain/spinal cord) and *NKX2*.*1* (ventral forebrain) were detected at very low levels in both control and SETBP1-/- cultures (2-ΔΔCT>30, data not shown). These observations suggest that neural induction and cortical fate specification occurred normally in the absence of SETBP1.

**Fig. 2.**
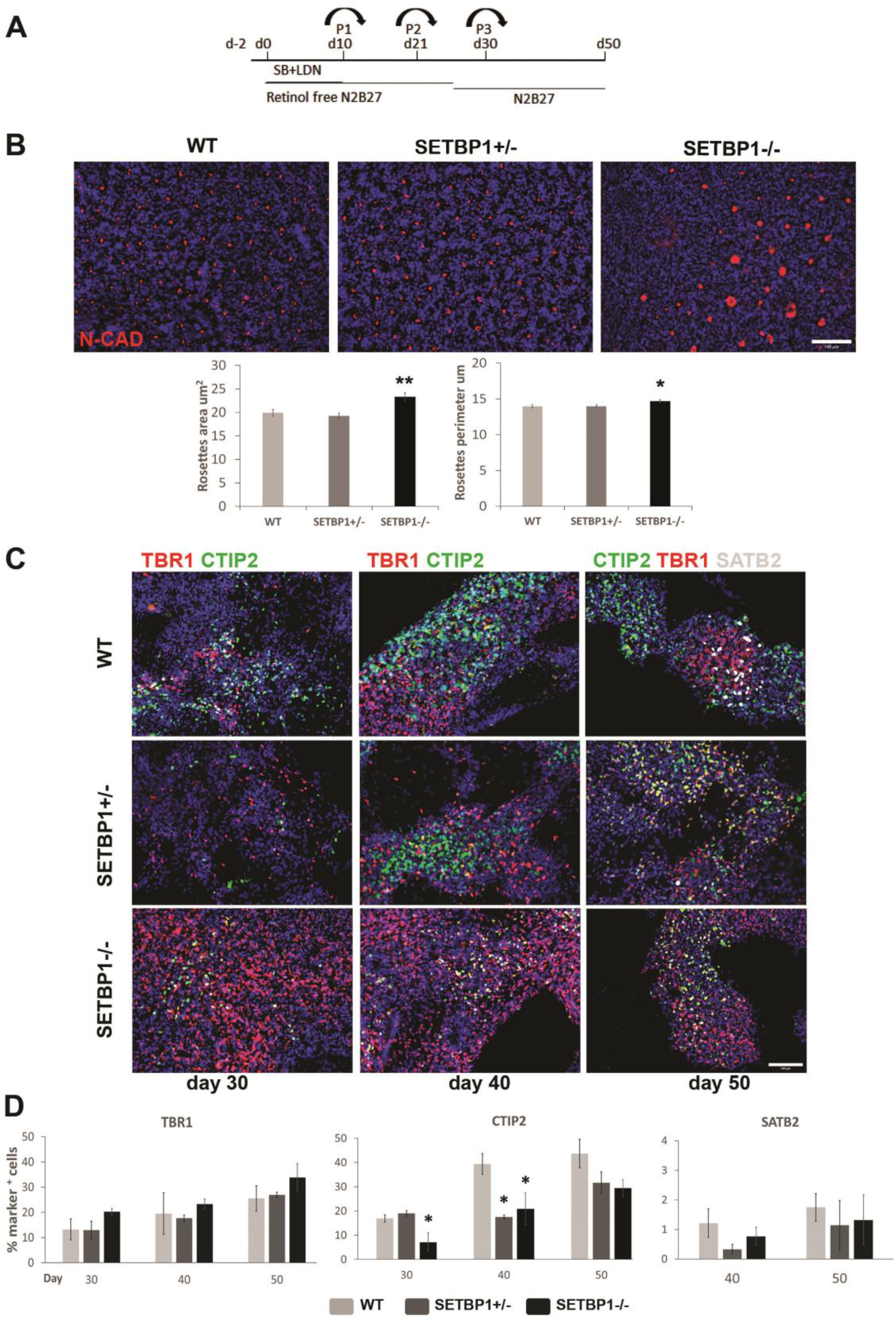
Cortical neuronal differentiation is impaired in SETBP1-deficient NPCs. (**A**) Schematic representation of hESC cortical differentiation protocol. (**B**) Day 18/20 cultures were immunostained for N-cadherin (red) and DAPI (blue) showing the organization and size of the neural rosettes. Graphs showing quantitative measurements for rosette apical area and perimeter (in uM) of a minimum of 900 rosettes per cell line (WT, HET1, HET2, KO1 and KO2) and ≥3000 rosettes per genotype WT, SETBP1+/- and SETBP1-/- (Mann-Whitney U test P=0.005 for rosettes area and P=0.025 for rosettes perimeter). (**C**) Immunostaining of cortical layer markers TBR1 (layer VI) and CTIP2 (layers V-VI) days 30, 40 and 50, respectively, and SATB2 (layers II-III) at day 50. Images representative of several independent experiments for each genotype (**D**) Quantitative analysis of the above. Data presented as mean ± s.e.m for each genotype with a minimum of two independent experiments carried out per line (WT = 5, HET1 and HET2 = 2, KO1 = 3, KO2 and KO3 = 2). One-way ANOVA test, Bonferroni Post Hoc (* p≤0.05, ** p≤0.01, *** p≤0.001). Scale bar: 100uM.

However, SETBP1-/- NPCs formed larger neural rosettes than that of the SETBP1+/- and WT cells (Fig. 2B). Neural rosettes are radial arrangements of polarized NPCs formed at around two-three weeks of hESC differentiation. Apart from the increased size (p=0.005 for rosettes area and p=0.025 for rosettes perimeter), SETBP1-/- rosettes, as visualised by N-CAD antibody staining, appear otherwise normal in terms of the shape and cellular organization.

### Cortical neuronal differentiation is altered in SETBP1-deficient NPCs

During development, corticogenesis is tightly regulated to ensure the generation of correct numbers of neurons in time and space. Neurons in different cortical layers are generated in an inside-out fashion, with neurons in the deep layers born first and upper layer neurons later(*26, 27*). Cortical differentiation of hESCs in vitro recapitulates the temporal aspects of this process(*28, 29*). We therefore performed immunostaining for two deep layer markers TBR1 (layer VI) and CTIP2 (BCL11B, layers V-VI) and an upper layer marker SATB2 (layers II and III) at day 30, 40 and 50, a time window that these neurons are being produced in the control cultures. We observed a decreased production of CTIP2^+^ cells in SETBP1-/- cultures compared to the WT controls at all time points (ANOVA day 30 P=0.030, day 40 P=0.037, day 50 P=0.214) (Fig. 2C and Fig. 2D). A reduction in CTIP2^+^ cells were also observed in SETBP1+/-cultures although statistics significance was only reached at day 40. Intriguingly, the opposite trend was observed for TBR1^+^ neurons that were over-represented in the SETBP1-/- cultures, although statistics significance was not reached (Fig. 2C, Fig. 2D and Fig. S3B). Consistent with being late born cells, SATB2^+^ neurons were not detected at day 30 and were very low in number even at day 40 and 50 (∼2%) in all cultures (Fig. 2C and Fig. 2D).

To investigate whether the observed differences in TBR1^+^ and CTIP2^+^ cells in SETBP1 mutant cultures was caused by a decrease in general neuronal production, we determined the total number of NPCs by immunostaining for PAX6 and NeuN neuronal marker at day 30, 40 and 50 (Fig. S4A and Fig. S4B). At all three time points SETBP1-/- cultures had a lower number of NeuN^+^ cells than the controls. Lower number of NeuN^+^ cells were also detected in SETBP1+/-cultures although no statistic differences. In contrast, while a large proportion of the WT cells no longer expressed progenitor marker PAX6 at day 40 (17% PAX6^+^ cells), around 45% cells in the SETBP1-/- cultures remained PAX6^+^ (Fig. S4A and Fig. S4B). Together, these findings demonstrate a distorted production of deep layer cortical neurons and an overall decrease in neuronal differentiation of cortical NPCs in SETBP1-/- cultures, while heterozygous deletion of SETBP1 had a milder effect on cortical differentiation in our model.

### SETBP1 deficiency results in increased cortical progenitor proliferation

The reduction of SETBP1-/- neurons could be due to an imbalance between NPC proliferation versus terminal differentiation. We therefore investigated a potential change in cell cycle of SETBP1-/- NPCs by EdU and Ki67 double labelling at day 34 (Fig. 3A-C). EdU is a thymidine analogue hence its incorporation marks cells in the S phase, while Ki67 is a protein present during all active phases of the cell cycle (G1, S, G2 and mitosis). The SETBP1+/- and WT cultures contained a similar number of EdU^+^ and Ki67^+^ cells (14.24 ± 1.10% vs 11.42 ± 0.11 % and 19.94 ± 1.21% vs 18.52 ± 0.93 %, respectively). However, significantly more EdU^+^ (28.27 ± 0.22%), Ki67^+^ (25.45 ± 1.14%) and EdU^+^Ki67^+^ cells (18.28 ± 0.56 % vs 12.3 ± 1.7%) were detected in the SETBP1-/- cultures (Fig. 3A-B). The fraction of EdU^+^Ki67^-^ cells within the EdU^+^ population is often used as an index for cell cycle exit(*30*), we found that the ratio of EdU^+^Ki67^-^/ EdU^+^ is lower in SETBP1-/- cultures than the SETBP1+/- and WT controls with a borderline p-value (P=0.055) (Fig. 3C), suggesting that SETBP1-deficient NPCs were slow in exiting the cell cycle compared to their isogenic counterparts. The ratio of EdU^+^Ki67^+^/ Ki67^+^ is inversely related to the length of cell cycle(*31*). Consistent with an increase in proliferation, this ratio was slightly higher in SETBP1-/- cultures than the controls, indicating the former have shorter cell cycle (Fig. 3C). To gain further insight into changes in cell cycle profile, we performed a flow cytometry based cell cycle analysis (Fig. 3D). This assay identifies cells in three major phases of the cell cycle (G0/1, S and G2/M) based on their DNA content. Since the cellular defects observed were largely limited to the SETBP1-/- cultures, we focused on this genotype in the subsequent studies. Compared to the WT control, the SETBP1-/- cultures contained a higher percentage of cells in S (9.24 ± 0.69 % vs 7.92 ± 1.52%) and G2/M phases (19.81 ± 0.18 % vs 18.07 ± 0.37%, P=0.029) and fewer cells in G0/G1 (68.01 ± 0.31% vs 70.24 ± 1.29 %). We also determined the number of cells in mitosis by antibody staining for phosphorylated histone H3 (PH3), no difference was observed between the SETBP1-/- and control cultures (Fig. 3A).

**Fig. 3.**
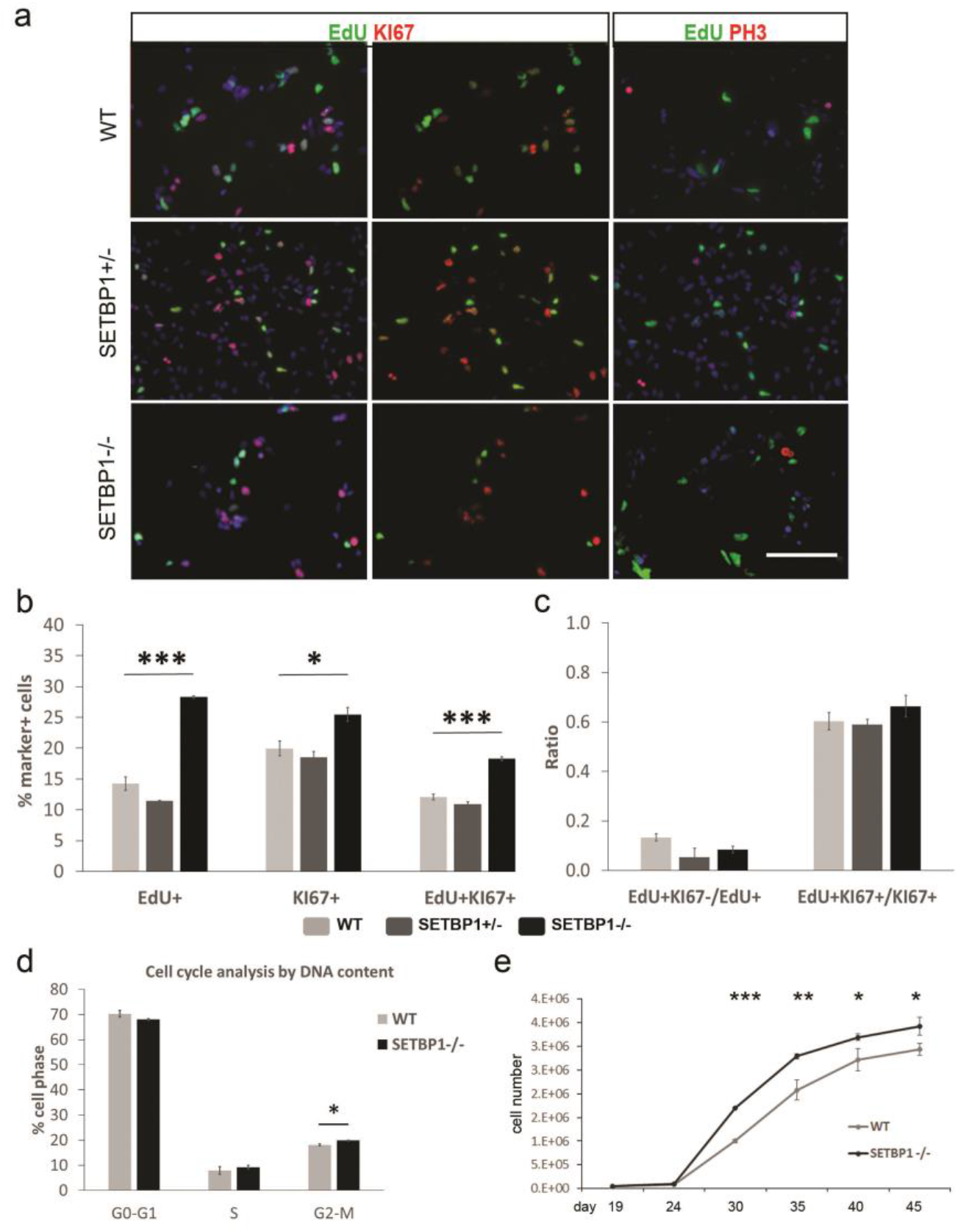
SETBP1 deficiency enhances cortical progenitor proliferation. (**A**) WT, SETBP1+/-(HET1) and SETBP1-/- (KO1) day 35 cultures were immunostained for EdU (green), Ki67 (red), PH3 (red) and counterstained with DAPI (blue). (**B**) Percentage of cells positive for EdU (P≤0.001), Ki67 (P=0.016) and EdU and Ki67 (cell cycle re-entry P≤0.001). (**C**) Ratios of cell cycle exit and cell cycle length (P=0.055, 0.043). Data presented as mean ± s.e.m from 3 independent wells with 6 random fields each. One-way ANOVA test (* p≤0.05, ** p≤0.01, *** p≤0.001). Scale bar: 100uM. (**D**) Cell cycle analysis by DNA content using Flow cytometry, % of cells in each of the cell cycle phases. Data presented as mean ± s.e.m of 2 independent experiments in triplicates. Student’s T test, one tail (G1-G0 P=0.118, S P=0.255, G2-M P=0.029). (**E**) Growth curve analysis from day 19 to day 45 showing the increased population growth of the SETBP1-/- (KO1) NPCs compared to the isogenic controls. Statistical significant differences were found from day 30 onwards (Student’s T test, P=6.71E-05 for day 30, 0.015 for day35, 0.033 for day 40, and 0.050 for day 45). Data presented as mean ± s.e.m from 3 independent wells with two technical measurements.

To investigate how altered cell cycle impact on the growth rate over time, we compared population growth of SETBP1-/- and WT cultures between day 19 and day 45. Consistent with EdU incorporation and cell cycle analysis, more cells were found in the SETBP1-/- cultures than the WT from day 30 onwards (P≤0.05, Fig. 3E). Together, these findings demonstrate that SETBP1 deficiency lead to enhanced NPC division by regulating cell cycle.

### Genome-wide transcriptome analysis identified Wnt/β-catenin signaling as a target of SETBP1 function

To gain further insight into the molecular mechanisms underlying prolonged proliferation window of SETBP1-deficient NPCs, we performed a genome wide transcriptome analysis of neural cells derived from the SETBP1-/- and isogenic WT control lines by RNA sequencing (RNAseq). To cover all stages of cellular abnormality, samples were collected from day 15 and day 21 (early and peak neural rosette stage, respectively) and day 34, when abnormal NPC division and neurogenesis was becoming evident. Principle Component Analysis (PCA) showed that 100% of the variance is attributed to SETBP1 genotypes, while the biological replicates within SETBP1-/- or the control samples exhibit 0% variance statistically (Fig. 4A). At a significant level of adjusted P ≤0.1, we identified 6060, 9997 and 17654 differentially expressed transcripts at the three analysed time points, respectively (Fig. 4B). Amongst the top differentially expressed genes, *FOXG1* was down regulated at day 15 and 21 (Fig. 4C-D), providing independent support of the previous observations by immunostaining (Fig. S2A-B) and RT-PCR (Fig. S3A). Other top down-regulated transcripts were several cortical transcription factors such as *LHX2, DMRTA2, SIX3, PAX6* and *EMX1*, with the changes most evident at day 15 and some recovery of transcription levels at day 21 (Fig. 4C-D). On the other hand, some cortical transcription factors such as *NESTIN, SOX1, HES1, EMX2 and OTX1/2* were up-regulated in the SETBP1-deficient NPCs at one or both of time points (Fig. 4C-D). No changes in expression level were observed for ventral telencephalic determinants such as *NKX2*.*1, GSX2* and *LHX6*, although these transcripts were present at very low abundance. At day 34, basal progenitor (*TBR2*), pan neuronal (*TUBB3, MAP2*) and cortical layers specific marker genes (*TBR1, CTIP2*/*BCL11B, CUX1/2, SATB2, RELN*) were found down-regulated in the SETBP1-deficient cultures. In contrast, *NESTIN* was 2-fold higher in the SETBP1-/- samples than the controls (Fig. 4E). These findings support the observed NPC SETBP1-deficient phenotype and demonstrate a further role for SETBP1 in cortical NPC proliferation and neurogenesis.

**Fig. 4.**
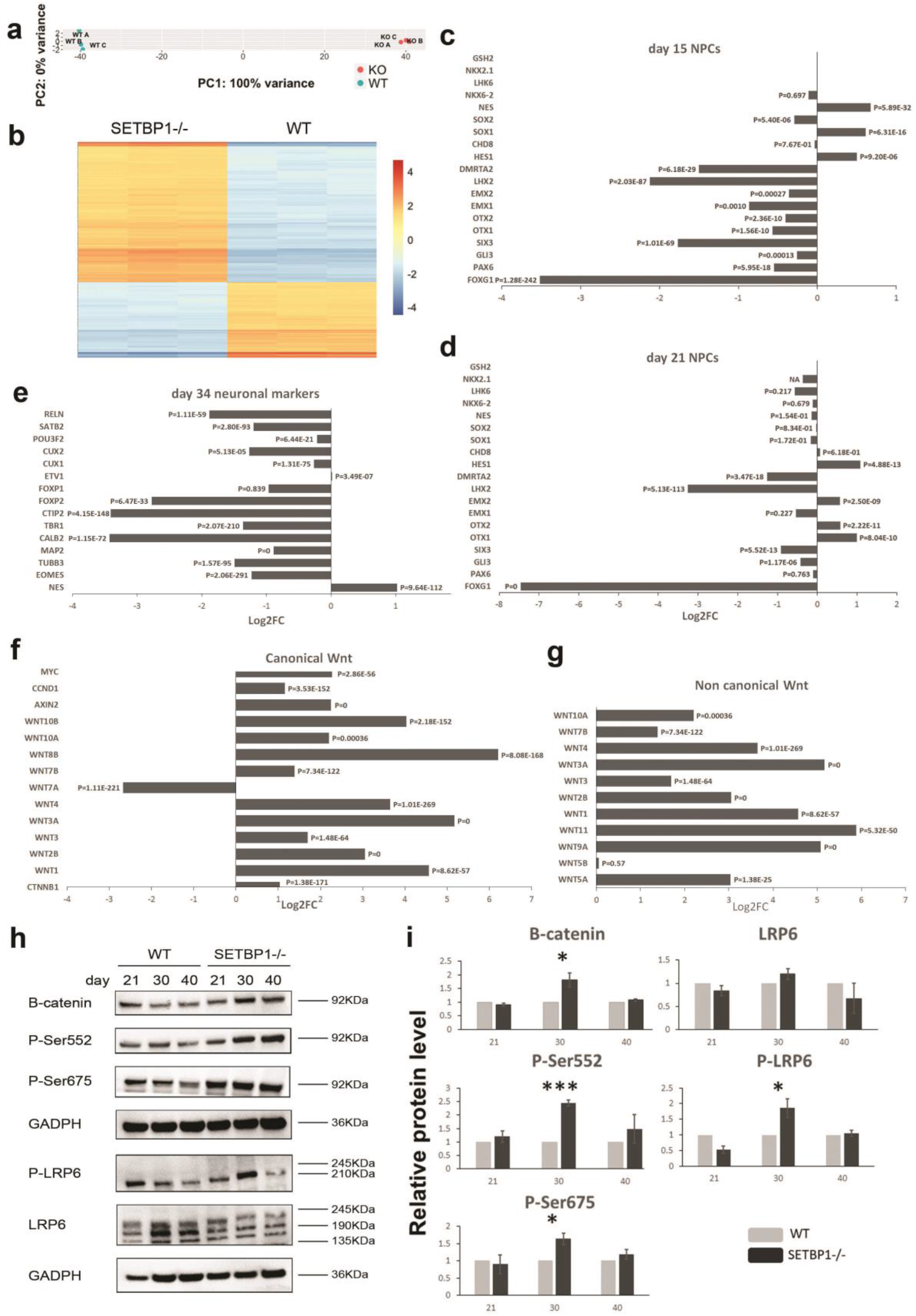
Genome wide transcriptome profiling revealed SETBP1 regulation of WNT signaling. (**A**) Principle component analysis (PCA) of the samples. (**B**) Heatmap depicting 17654 differentially expressed transcripts at day 34 (padj ≤ 0.1). (**C-E**) Example of differentially expressed genes associated with telencephalic patterning at day 15, day 21 and neuron genes at day 34, respectively. (**F-G**) Differentially expressed genes associated with canonical non-canonical Wnt pathway at day 34. (**H**) Representative images of Western blot analysis for WNT signaling proteins for WT and SETBP1-/- (KO1). (**I**) Relative protein level of β-catenin, β-catenin p-S552 and p-S675, and LRP6 co-receptor and p-LRP6 at day 21, 30 and 40 relative to WT basal levels. Data from 3 independent differentiations analysed in duplicates or triplicates. Student’s T test was used to compare the expression between the two lines. (* p ≤ 0.05, **p ≤ 0.01, ***p ≤ 0.001).

Using DAVID 6.8 gene functional classification tool (https://david.ncifcrf.gov/) on the top 1000 differentially expressed protein coding genes, we identified that top enriched gene ontology (GO) terms concerned mainly biological processes such as regulation of transcription, cell adhesion and extracellular matrix organization. KEGG (Kyoto Encyclopedia of Genes and Genomes) pathway analysis revealed DNA replication, Wnt, hippo, PI3K-Akt and ECM-receptor interaction signaling as the top enriched pathways up-regulated in SETBP1-/- cultures (Table S2-4). All these pathways are highly relevant to the regulation of NPCs proliferation and neurogenesis (*32-36*).

Wnt signaling is known to play an important role in cortical development. Altered Wnt pathway was identified at all three differentiation stages, with the biggest changes observed at day 21 (FC 2.73, Padj=0.0029) and day 34 (FC 2.53, Padj=0.0014, Table S3-4). We therefore examined the gene set for genes involved in Wnt signaling (Fig. 4F-G and Figure S3). Strikingly, the majority of the WNT ligands, both canonical and non-canonical, were highly up-regulated in SETBP1-deficient cells, with FC varying from 2.5 to 73. Also up-regulated were the canonical WNT/β-catenin signaling responsive genes *C-MYC* (4.8x), *CYCLIND1* (*CCND1*, 2x) and *AXIN2* (4.7x, Fig. 4F). In contrast, genes involved in β-catenin degradation complex (*GSK3* β, *CSNK, AXIN1/2, APC*) were mostly down-regulated (Fig. S5).

To ascertain that WNT/β-catenin signaling is indeed elevated in SETBP1-deficient NPCs at the protein level, we determined the level of β-catenin and WNT co-receptor LRP6 in day 21, day 30 and day 40 neural cultures by Western blot. Activation of the canonical WNT signaling results in N-terminal phosphorylation of β-catenin by GSK3β, leading to degradation of β-catenin(*37, 38*). We found that the level of total β-catenin was significantly higher in SETBP1-/- cultures than the controls at day 30 (P=0.031), although no differences were found at day 21 and 40. It has been reported previously that C-terminal phosphorylation of β-catenin in serine 552 and serine 675 (p-S552 and p-S675) by AKT and PKA can enhance β-catenin/TCF reporter activation(*39, 40*). We detected an average of 2.5-fold increase of p-S552 (p=0.00024) and 1.5 fold increase of p-S675 (p=0.019) in SETBP1-/- cultures than the controls at day 30 (Fig. 4H-I).

Another key phosphorylation event in the activation of the WNT signaling cascade is the phosphorylation of the LRP5 and LRP6 co-receptors(*41, 42*), LRP6 is known to play a more dominant role during embryogenesis. We observed a near 2-fold increase of phosphorylated LRP6 (p-LRP6) in day 30 SETBP1-/- NPCs than the isogenic control cells (p=0.046, Fig. 4H-I). Together, these studies validated the increase of Wnt/β-catenin activation in SETBP1-deficient cells and provide the first demonstration of a regulatory role for SETBP1 in canonical WNT signaling in cortical NPCs.

### Pharmacological inhibition of Wnt/β-catenin pathway rescues proliferation defect of SETBP1-/- NPCs

To establish a causal relationship between the increased Wnt/β-catenin signaling and over proliferation of SETBP1-deficient NPCs, we interrogated Wnt signaling using XAV939 (XAV), a small molecule tankyrase inhibitor that stabilizes Axin and stimulates β-catenin degradation(*43*). SETBP1-/-, SETBP1+/- and WT NPC cultures were exposed to XAV for 10 days from day 11, a time window prior to the phenotypic manifestation (Fig. 5A and Figure S6A). Wnt signaling inhibition by XAV was verified by evident reduction in total β-catenin, p-S552/p-S675 as well as p-LRP6 comparing treated SETBP1-/- with respective to the no XAV sister cultures in both WT and SETBP1-/- cultures (Fig. 5B). Importantly, after XAV treatment, total β-catenin, p-S552 and p-S675 in SETBP1-/- cells were no longer different to the isogenic control cells without XAV treatment. As a control for inhibitor specificity, the levels of the GAPDH were not affected by XAV treatment.

**Fig. 5.**
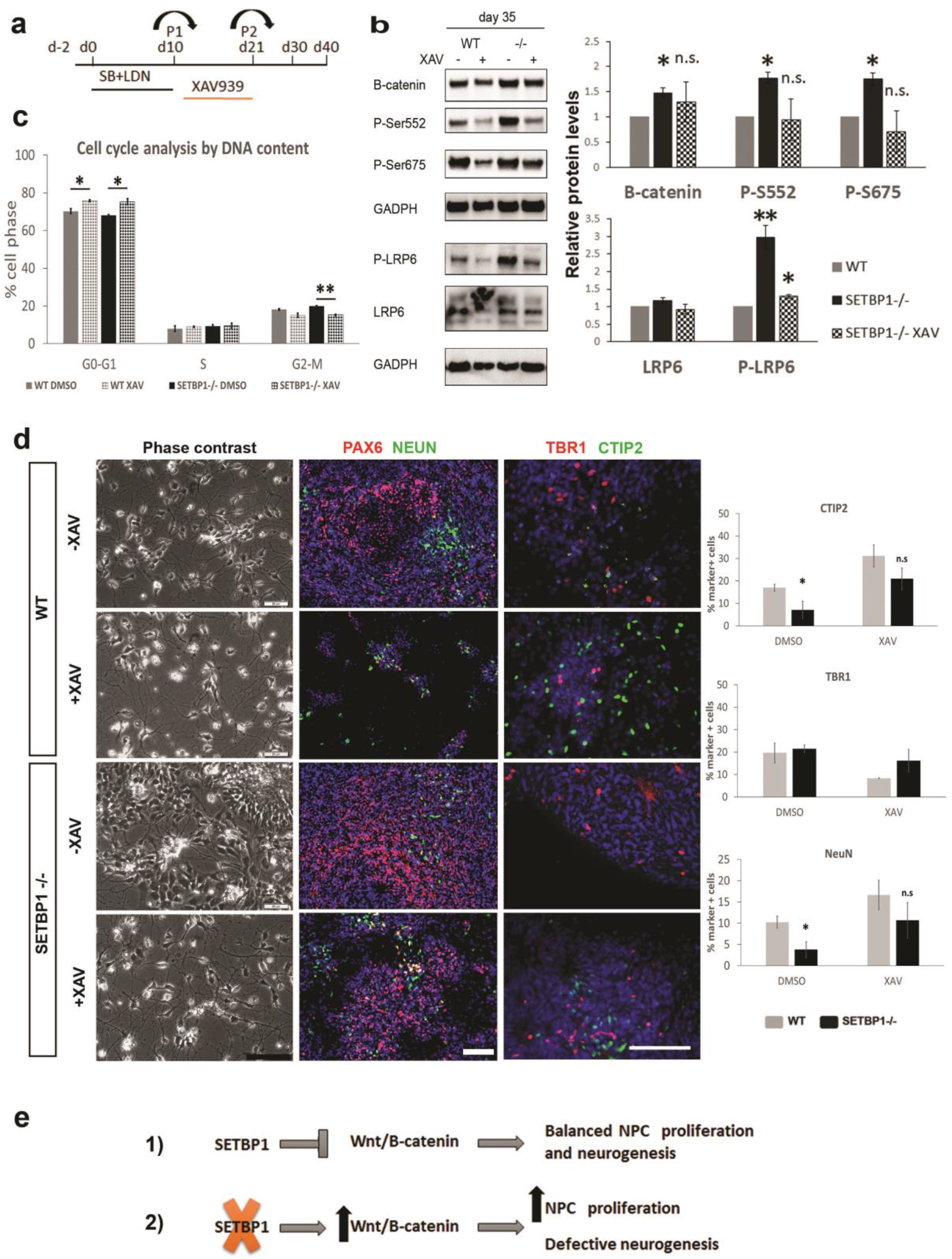
Functional interrogation of WNT signaling with XAV939 treatment. (**A**) Experimental scheme. Differentiation cultures under basal condition (control) or exposed to XAV939 2uM from day 11 to day 21. (**B**) Western blot analysis for the effects of XAV treatment on WNT signaling. SETBP1-/- (KO1) protein levels after XAV treatment at day 35 relative to WT basal levels. Data from 2 independent differentiations analysed in duplicates or triplicates. Student’s T test was used to compare the expression between the two lines, β-catenin basal P=0.033, XAV P=0.542, S552 basal P=0.023, XAV P=0.906, S674 basal P=0.024, XAV P=0.554, LRP6 basal P=0.153, XAV P=0.644, P-LRP6 basal P=0.004, XAV P=0.012. (**C**) Analysis of the effect of XAV treatment on the proportion of cells in each of the cell cycle phases at day 35. Data presented as mean ± s.e.m of 2 independent experiments in triplicates. Student’s T test, one tail, was used to compare the expression between the two lines and the two conditions (WT basal vs XAV G0-G1 P=0.029, S P≥0.05, G2-M P≥0.05, SETBP1-/- basal vs XAV G0-G1 P=0.021, S P≥0.05, G2-M P=0.0045. (**D**) Phase contrast and immunofluorescence images of cultures in basal or XAV treated conditions at day 30. Cell nuclei were labelled by DAPI. Scale bar: 100uM. Bar-graphs showing quantification of CTIP2, TBR1 and NeuN positive neurons. Student’s T test was used to compare the expression between the two lines, CTIP2^+^ cells: basal P=0.029, XAV P=0.214; TBR1^+^ cells: basal P=0.672 XAV P=0.258, NeuN^+^ cells: basal P=0.038, XAV P=0.333 (**E**) Schematic illustration depicting the role of SETBP1 in regulating NPC proliferation and neurogenesis. 1) In normal conditions, adequate level of Wnt/β-catenin signaling is modulated by SETBP1 to ensure a balanced NPC proliferation and neurogenesis. 2) Loss of SETBP1 leads to elevated Wnt/β-catenin signaling leading to excessive NPC expansion and defective neurogenesis. (Student’s T test * p ≤ 0.05, **p ≤ 0.01, ***p ≤ 0.001).

The effect on XAV treatment in NPCs proliferation was examined via cell cycle analysis of DNA content (Fig. 5C). An increase of cells in G1 phase and a decrease of cells in G2-M was observed for both SETBP1-/- and control NPCs. In XAV treated SETBP1-/- cultures, the number of cells in G2-M phase was restored to a level similar to those in the WT cultures with or without XAV (15.18 ± 0.51 and 15.02 ± 1.07, P=0.902). We next examined the effect of XAV on NPC and neuronal numbers at day 30, 40 and 50 (Fig. 5D and Fig. S6B). Compared to non-treated SETBP1-/- cultures, cells in XAV treated cultures exhibited pronounced neuronal arborisation similarly to those in the WT control cultures without XAV (Fig. 5D and Fig. S6B). Immunostaining revealed that, as expected, XAV treatment in the WT cultures accelerated neuronal differentiation as demonstrated by a reduced number of PAX6^+^ and NESTIN^+^ cells and concurrent increase of NeuN^+^ and MAP2^+^ cells in comparison to no XAV sister control cultures (Fig. 5D and Fig. S6B). XAV treatment also resulted in a significant increase in CTIP2^+^ cells and was accompanied by a reduction in TBR1^+^ cells from day 30 onwards and higher detection of SATB2^+^ cells (Fig. 5D and Fig. S6B).

Together, our data demonstrates that inhibition of WNT/β-catenin signaling can restore the proliferation and neurogenesis defects of SETBP1-/- NPCs and thus provide a functional verification that SETBP1 is playing a role in WNT signaling.

## DISCUSSION

The generation of neurons in the cortex is tightly regulated temporally and spatially with a progressive temporal restriction in progenitor potential. During the course of cortical development, NPCs firstly reproduce themselves to expand the population of progenitors via symmetric or proliferative cell divisions(*44, 45*). Later, cell division pattern changes to asymmetric neurogenic divisions that generates an NPC that re-enters the cell cycle and a postmitotic neuron, or symmetric neurogenic divisions that yield two neurons(*26, 46*). Defects in this process can lead to a wide range of brain malformations such as micro- or macrocephaly. Using genome edited hESC and in vitro cortical differentiation as an experimental model, we report here that loss of SETBP1 resulted in protracted NPC proliferation and defective neurogenesis, thus identifying SETBP1 as an important regulator governing the delicate balance between NPC expansion and terminal differentiation. This newly discovered biological function of SETBP1 in human neural development is consistent with its high-level expression in the developing cortex and its evolutionary conservation.

Dysregulation of WNT signaling in SETBP1-deficient neuronal cultures presents another interesting new finding of this study. This regulatory relationship was demonstrated at both transcript and protein level with further support of functional interrogation and phenotypic rescue. WNT signaling is known to play an important role in cortical development. Elevated canonical WNT signaling by enforced expression of stabilized β-catenin promotes cell cycle re-entry of NPCs, leading to their excessive expansion in telencephalon(*47*). A prominent feature of our SETBP1-deficiency model is prolonged NPC proliferation, due to shortened cell-cycle length and reduced cell cycle exit rate. Therefore, the increased WNT signaling in SETBP1-/- cultures is likely a key contributor to the aberrant NPC proliferation.

Interestingly, dysregulated WNT signaling have been recently reported in a growing number of studies employing patient-derived iPSC or CRISPR/CAS9 edited hESC models of neurodevelopmental disorders that include schizophrenia, autism spectrum disorder (ASD) and intellectual disability(*48-52*). Elevated WNT signaling has been implicated as a cause of the macrocephaly observed in ASD patients(*53*). The current study suggests that increased WNT signaling may be also an underlying mechanism of the cognitive and motor impairment observed in patients with SETBP1 disorder.

SETBP1 is well known as an inhibitor of PP2A in acute myeloid leukemia (*54*). A transcription factor function was first described in murine myeloid progenitor cells through binding of Hoxa9/10 promoters(*55*). Using a similar model, the same group described that binding of Setbp1 to Runx1 promoter caused a downregulation of RUNX1 expression (*56*). In a recent study, Piazza and colleagues show the ability of SETBP1 to bind to gDNA in AT-rich promoter regions triggering activation of gene expression via the recruitment of HCF1/KMT2A/PHF8 epigenetic complex (*57*). They also described that in utero electroporation of SETBP1-G870S in the developing mouse brain caused an impairment in neurogenesis and delay in neuronal migration. A recent paper modelling SGS using iPSCs show that SGS NPCs present aberrant proliferation, DNA damage and dysregulated pathways related with cancer and apoptosis(*58*). However, these studies were carried mostly in myeloid progenitor cells or in SGS mutation related models that resemble SETBP1 gain of function. A role of SETBP1 directly regulating WNT signaling in the context of human brain has not yet been reported. While further investigation is needed to unravel the role of SETBP1 in WNT/β-catenin signaling in the context of cortical development, our work shows a new pathway to study in the context of SETBP1 haploinsufficiency and a possible therapeutic road to explore.

*FOXG1* is the most significantly down-regulated transcript in SETBP1-/- NPC cultures. Foxg1 has been shown to suppress Wnt signaling in several mouse models(*59, 60*). Therefore, reduced FOXG1 may contribute to the elevated WNT activity in SETBP1-deficient cultures. SETBP1 deficiency resulted in a significant reduction of CTIP2^+^ cells and relatively higher numbers of TBR1^+^ cells. Previous studies in rodents suggest that Foxg1 confers the competence of cortical progenitors for the characteristic ordered generation of layer-specific neuronal subtypes by coordinating Wnt and Shh signaling pathways in the telencephalon (*61, 62*). Therefore, we cannot exclude the possibility that the distorted neuronal production in our SETBP1-/- cultures may be attributed at least in part to FOXG1 hypo function. Our data opens new avenues to explore the functional link between FOXG1 and SETBP1 genes in the context of brain development.

Disturbed NPC proliferation and neuronal differentiation can lead to brain malformation as it happens in epilepsy with heteropia, ASD microcephaly or lissencephaly patients(*4-7*). Our finding of disturbed cortical progenitor proliferation and defective neurogenesis in SETBP1-/- model lend itself an invaluable tool to further investigate the aetiology of SETBP1 disorder and may contribute to the search of therapeutic compounds.

## MATERIALS AND METHODS

### HESC culture and cortical neural differentiation

HESCs (H7-WA07) and genome edited H7 derivatives were maintained on Matrigel-coated plates in Essential 8 media (TeSR-E8, Stemcell Technologies)(*25*). All hESCs were passaged via manual dissociation using Gentle Cell Dissociation Reagent (Stemcell Technologies). For differentiation, hESCs were pre-plated on growth factor-reduced matrigel in TeSR-E8. When cells reached >80% confluence, neural differentiation was initiated by switching TeSR-E8 to DMEM-F12/Neurobasal (2:1) supplemented with N2 and B27 (referred to thereafter as N2B27). For the first 10 days, cultures were supplemented with SB431542 (10μM, Tocris) and LDN-193189 (100nM, StemGene). Cultures were passaged using EDTA firstly on day 10 at a ratio of 1:2 onto fibronectin-coated plates. The second and third split were performed on day 20-21 and day 30, respectively, onto poly-D-lysine/laminin coated 24-well plates at a density of 125.000 cells/well. Retinol-free B27 was used for the first 25 days, followed by normal B27 from day 26 onwards. For Wnt/B-catenin inhibition, XAV939 (2μM, SelleckChem) was added to the media for 10 days after the second split.

### CRISPR/CAS9 genome editing

Guide RNAs (gRNAs) were designed to target exon 4 of the human *SETBP1* gene using two independent CRISPR gRNA design tools: Atum CRISPR gRNA (former DNA2.0 https://www.atum.bio/eCommerce/cas9/input) and the CRISPR Design Tool (http://crispr.mit.edu) to minimize the risk of off-target effects of Cas9 nuclease. gRNA1 5′-TGTGGCCGGCTTCGCTGTGCTGG, gRNA2 5′-GGAGGTCATCGCGGTTTTGCAGG gRNA3 5’-TGAAATTTCATCTCGCTCATGGG. All gRNAs were synthesized as oligonucleotides and cloned into the pSpCas9(BB)-2A-GFP plasmid (px458, Addgene) following the protocol of Ran et al(*23*). A donor template (gene targeting) vector for homologous recombination was constructed that contains a PGK-puro-pA selection cassette flanked by a 502 bp 5’ homologous arm corresponding to part of exon 4 and 551 bp 3’ homologous arm in intron 4/5. All three gRNA target sites are located within the 2705bp region between the two homologous arms (Figure 1). HESCs were transfected with a total of 4ug DNA in a ratio 2:3 (gRNAs:donor template) using the Amaxa P3 Primary Cell 4D-Nucleofector Kit (Lonza). Puromycin was added 3 days after electroporation at a concentration of 0.5ug/ml, drug resistant hESC colonies were picked one week later and expanded clonally. Genotyping was done by genomic PCR (primers used are provided in Table S1) followed by Sanger sequencing of candidate mutant PCR product. Only SETBP1 heterozygous lines were generated in the first round of transfection (Fig. 1B lanes 2, 3 and 7 for mutant clones confirmed with 5’ and 3’ PCRs). One of the heterozygous was used for a second round of targeting using the same gRNAs without the donor plasmid, which yield several clones carrying the original KO allele and a 5bp deletion in the other allele (Fig. S1B-C). Potential off-target effects were analyzed by genomic PCR followed by Sanger sequencing of the PCR product. No disruption of the WT sequence was detected for any of the tested cell lines (data not shown).

### Karyotyping

Sub-confluent hESCs cultures were treated with 0.1μg/ml Demecolcine (Sigma D1925) for 1 hour at 37°C and then dissociated to a single cell suspension using Accutase (ThermoFisher) for 10 minutes at 37°C. Cells were collected and washed twice with PBS by centrifugation for 4 min at 900 rpm. Cells were resuspended in 2 ml of PBS and 6 ml of 0.075 M KCl hypotonic solution was added to the tubes following incubation at 37°C 15 min. Additional 4 ml of 0.075 KCl was added after incubation and cells were collected by centrifugation for 4 min at 900 rpm. The supernatant was removed leaving 300 μl to resuspend the cell pellet by flicking. 4 ml of pre-chilled (−20°C) methanol/acetic acid (3:1, VWR chemicals) was added dropwise and flicking to homogenize. Cell suspension was incubated for 30 min at room temperature. Cells were then centrifuged for 4 minutes at 900 rpm and resuspended with additional 4 ml of methanol/acetic acid. Cells were collected as previously and resuspended in 300 μl of methanol/acetic acid. Cell suspension was dropped onto a slide (pre-chilled and laid on angle) from a height of around 30 cm. Slides were air dried and chromosome spread was stained and mounted using a mix of mounting media with DAPI (1:3000). Images of chromosome spreads were obtained with an inverted microscope. Images were acquired at 100x using the Leica Application Suite software and manually counted on ImageJ.

### Immunocytochemistry and EdU-labelling

Cultures were fixed with 4% (w/v) paraformaldehyde and permeabilised with 0.1% (v/v) Triton X-100 in PBS. Following blocking with 1% (w/v) bovine serum albumin and 3% (v/v) donkey serum, cells were incubated with primary antibodies overnight at 4°C. After three washes with PBS, cells were incubated with complementary Alexa Fluor-conjugated antibodies and counterstained with DAPI. All antibodies were diluted in PBS-T 1% donkey serum and incubated overnight at 4 degrees. Secondary antibodies were diluted (1:1000, Life technologies) in PBS-T 1% donkey serum and incubated for 1 h at room temperature. The primary antibodies used are: Goat anti-OCT4, 1/500 (Santa Cruz), Goat anti-SOX2, 1/200 (Santa Cruz), Mouse anti-TRA1-60 and Mouse anti-TRA1-81 1/200 (both Millipore), Mouse anti-PAX6, 1/1000 (DHSB), Rabbit anti-OTX2, 1/300, Rabbit anti-NEUN, 1/500 (both Millipore), Mouse anti-NESTIN, 1/300 (BD), Mouse anti-N-CAD, 1/100 (Life technologies), Mouse anti-ki67, 1/1000 (Leica biosystems), Rabbit anti-PH3, 1/1000, Rabbit anti-TBR1, 1/500, Rat anti-CTIP2, 1/500, Mouse anti-SATB2, 1/50 (all from Abcam).

To measure cell proliferation, cultures were incubated with 10 μM EdU (5-ethynyl-2’-deoxyuridine) for 30 min before fixation in the case of hESC cultures and 2 h for NPC cultures. EdU detection was carried out using the Click-iT EdU Alexa Fluor 488/555 imaging kit as per manufacturer instructions (Life Technologies).

Images were acquired using a DMI600b inverted microscope (Leica Microsystems). Cell counting was carried out using the CellProfiler or FIJI (Nucleus counter or cell counter plugins) software to analyse a minimum of 5-10 randomly placed fields of view per stain. Data were collected from two to five independent differentiation runs, sample size (n) per experiment and genotype indicated in the Figure legends.

### Quantitative RT-PCR (qPCR)

Total RNA was extracted using TRIzol (Invitrogen) and treated with TURBO DNA-free (Ambion). cDNA was generated using qScript cDNA synthesis kit (Quanta Biosciences). qPCR was performed with Mesa Green qPCR master mix (Eurogentec) with specific primers listed in (primers Supplementary Table 1). When possible, primers were designed to encompass exon-exon junctions. Cq values were normalised to *GAPDH* housekeeping reference gene and changes in expression level were calculated using the 2-ΔΔCT method(*63*). All data were obtained from 3 independent differentiations with PCR carried out in 2 independent runs each with three technical replicates. All PCRs were run on a QuantStudio Real-time PCR machine (Applied Biosystems).

### Flow cytometry

Cultured CNPs at differentiation day 34 were dissociated in Accutase (ThermoFisher) for 10 minutes at 37 °C, then washed in PBS and counted. Samples containing 3*10^6 cells were fixed in 70% EtOH at -20°C overnight. The cells were washed three times in DPBS and blocked in solution 1%BSA-3% donkey serum for 45 minutes. Following Mouse anti-NESTIN (1/300, BD) antibody in 1%BSA-1% donkey serum or mouse anti IgG, 2h at RT. Cells were washed in PBS three times and incubated in secondary antibody (alexa488 1:1000, Life technologies) in 1%BSA-1% donkey serum for 1h at RT. Cells were washed twice in PBS and incubated with RNaseA (200 μg/ml, ThermoFisher) for 30 minutes. Then centrifuged and treated with DAPI (0.3 μg/ml, ThermoFisher) for 10 minutes. Cells were washed twice in DPBS and resuspended in 0.5 ml to be filtrated to remove clumps (Corning™ Falcon™ Test Tube with Cell Strainer Snap Cap). The samples were analysed on a BD LSRFortessa cell analyser (BD Biosciences). Data was analyzed in FlowJo (BD Biosciences) and Statistical analysis was performed in SPSS (IBM).

### Growth curve study

Cells were seeded in triplicate at 50,000 cells/well onto poly-D-lysine/laminin coated 48 well plates at day 19. Retinol-free B27 was used for the first 25 days, followed by normal B27 from day 26 onwards. Cells were dissociated into single cells every 4-5 days and counted manually in a Neubauer chamber.

### RNAseq

RNA was extracted and purified using the PureLink RNA Mini Kit (Thermofisher Scientific). Libraries were prepared using the TruSeq Stranded mRNA kit (Illumina) from 1μg RNA extracted from 3 biological replicate samples each collected at 3 time points of differentiation (days 15, 21, and 34, n=9 each for SETBP1-/- and the isogenic control cells, respectively). 75bp paired-end sequencing was performed on a HiSeq 4000 sequencer (Illumina, USA) yielding 30 – 45 million reads per sample. Reads were mapped to the human genome (GRCh38) using Burrows-Wheeler Aligner algorithms(*64*) and individual gene read counts calculated using featureCounts(*65*). DeSeq2 was used to calculate differential gene expression with a cut-off of adjusted p-value<0.1, FDR of 10% and a FC>1.5 (*66*). Gene Ontology functional enrichment for biological processes was performed using DAVID (v6.8) for the top 1000 more significant genes (ranked for p adjusted and FC value), with all the protein coding genes in our dataset as background (*67*). Calculated p values were adjusted for multiple testing using the Benjamin-Hochberg correction. Raw sequence data files are publicly available from the NCBI Gene Expression Omnibus (GSE180185).

### Western Blotting

Protein extraction was performed with RIPA buffer (NEB) in the presence of protease and phosphatase inhibitors (Sigma) and quantified using the Bio-Rad DC protein assay (Bio-Rad). Total protein lysates (10-15 ug) were resolved in Bolt bistris plus 4-12% gels (Life technologies). PVDF membranes were blocked for 2h in 5% (w/v) BSA TBST buffer and the following primary antibodies diluted in blocking buffer were used: Mouse anti-GAPDH (1/5000, Abcam), mouse anti-B-catenin (sc7963, 1/1000, Santa Cruz Biotechnology), rabbit anti-P-Ser552 B-catenin (1/500, Cell signalling), rabbit anti-P-Ser675 B-catenin (1/500, Cell signalling), rabbit anti-LRP6 (1/500, Cell signalling), rabbit anti-P-LRP6 (1/500, Cell signalling). Incubation was performed overnight at 4C and primary antibodies were detected with anti-rabbit and anti-mouse HRP antibodies (Abcam) using the Luminata Crescendo Western HRP substrate (Millipore). Protein samples from 3 independent differentiations were analysed except the XAV treatment analysis where 2 rounds of differentiation were performed.

### Statistical Analysis

Statistical analyses were performed using IBM SPSS 23 software. Student’s T test or Mann-Withney U test were used for comparisons between two groups. One-way ANOVA and Kruskal-Wallis Test were used for comparisons between three groups Statistically significant differences were considered when p-value≤0.05. Two-tailed test was used unless indicated otherwise.

## Acknowledgements

We thank Dr. Robert Andrews and Dr. Daniel Cabezas de la Fuente for invaluable assistance with RNAseq data analysis. We acknowledge Ms. Eliza Wide contribution to the study during her MSc project. Thanks also to all members of the Li laboratory for helpful discussions during the course of this study. RNAseq was performed at the Oxford Genomics Centre.

This work was funded by the Medical Research Council of the United Kingdom grant 506390.

## Author Contributions

L.F.C. and M.L. conceived and designed the study. L.F.C. carried out and analysed the data. L.F.C. and M.L. interpreted the data and wrote the paper.

