## supplementary material for "WNT/β-catenin dependant alteration of cortical neurogenesis in a human stem cell model of SETBP1 disorder"

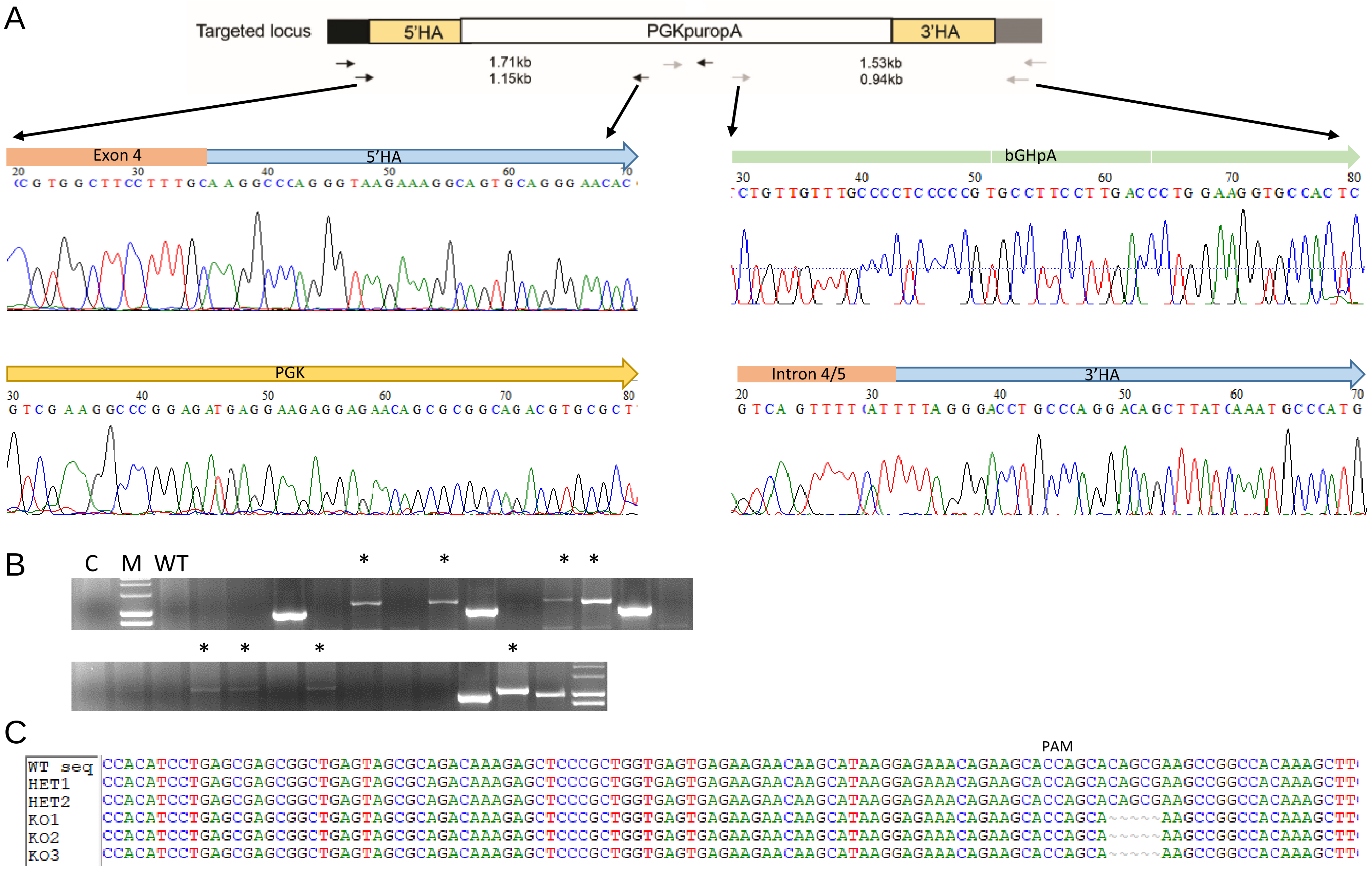
A
Targeted locus
5'HA
PGKpuropA
3'HA
1.71kb
1.15kb
1.53kb
0.94kb
Exon 4
5'HA
bGHpA
PGK
Intron 4/5
3'HA
B
C
M
WT
C
PAM
WT seq CCACATCCTGAGCGAGCGGCTGAGTAGCGCAGACAAAGAGCTCCCGCTGGTGAGTGAGAAGAACAAGCATAAGGAGAAACAGAAGCACCAGCACAGCGAAGCCGGCCACAAAGCTT
HET1 CCACATCCTGAGCGAGCGGCTGAGTAGCGCAGACAAAGAGCTCCCGCTGGTGAGTGAGAAGAACAAGCATAAGGAGAAACAGAAGCACCAGCACAGCGAAGCCGGCCACAAAGCTT
HET2 CCACATCCTGAGCGAGCGGCTGAGTAGCGCAGACAAAGAGCTCCCGCTGGTGAGTGAGAAGAACAAGCATAAGGAGAAACAGAAGCACCAGCACAGCGAAGCCGGCCACAAAGCTT
KO1 CCACATCCTGAGCGAGCGGCTGAGTAGCGCAGACAAAGAGCTCCCGCTGGTGAGTGAGAAGAACAAGCATAAGGAGAAACAGAAGCACCAGCA~~~~~AAGCCGGCCACAAAGCTT
KO2 CCACATCCTGAGCGAGCGGCTGAGTAGCGCAGACAAAGAGCTCCCGCTGGTGAGTGAGAAGAACAAGCATAAGGAGAAACAGAAGCACCAGCA~~~~~AAGCCGGCCACAAAGCTT
KO3 CCACATCCTGAGCGAGCGGCTGAGTAGCGCAGACAAAGAGCTCCCGCTGGTGAGTGAGAAGAACAAGCATAAGGAGAAACAGAAGCACCAGCA~~~~~AAGCCGGCCACAAAGCTT

**Fig. S1. Extra data regarding characterization of the SETBP1-deficient hESC lines**. (**A**) Representation of 5’ and 3’ PCR amplicon sequences. Diagram shows sequence reading from outside 5’HA (Forward sequence) into the PGK promoter (Reverse sequence) and from bGHpA (Forward sequence) towards the end of 3’HA and beginning of intronic sequence (Reverse sequence). (**B**) Agarose gels showing the 5’HA PCR amplicon from second round of targeting, * indicates positive clones. (**C**) Alignment of the cell lines used in the study for the sequence flanking gRNA1 target locus. A deletion of 5bp (CAGCG) was detected on the KO clones (SETBP1-/-).

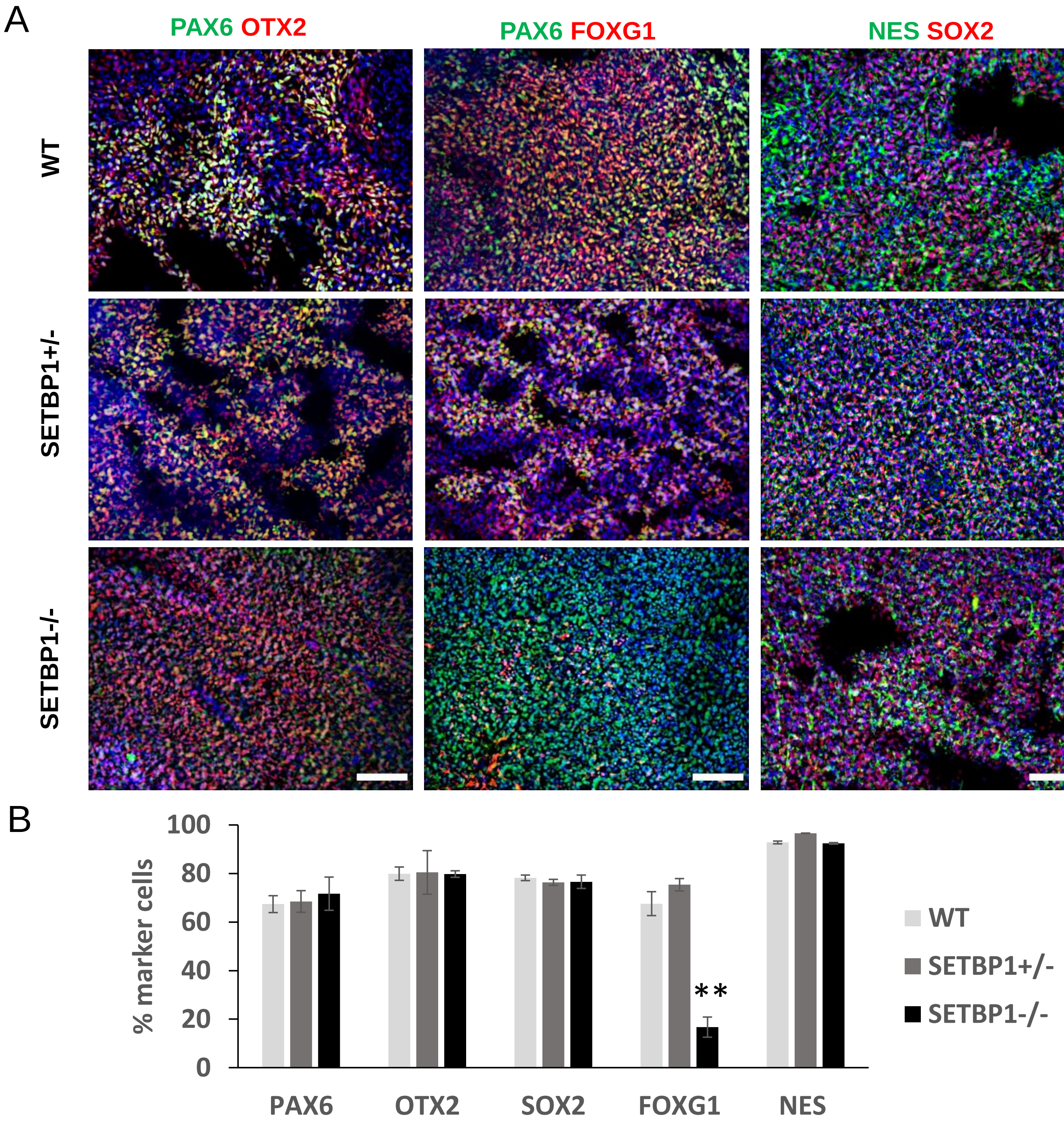
A
PAX6 OTX2
PAX6 FOXG1
NES SOX2
WT
SETBP1+/-
SETBP1-/-
B
% marker cells
100
80
60
40
20
0
PAX6
OTX2
SOX2
FOXG1
NES
**
WT
SETBP1+/-
SETBP1-/-

**Fig. S2. Expression of NPC markers at day 18-21.** (**A**) Day 18-21 cultures were immunostained for PAX6 (green), OTX2 (red), NES (green), SOX2 (red) and FOXG1 (red). Dapi was used to label all nuclei. Scale bar: 100uM. (**B**) Quantitative data presented as mean ± s.e.m for each genotype with a minimum of two independent experiments carried out per line and minimum of four per genotype (WT = 7, HET1 and HET2 = 2, KO1 = 6, KO2 = 4 and KO3 = 2). One-way ANOVA test or Kruskal-Wallis non-parametric test, PAX6 P=0.815, OTX2 P=1, SOX2 P=0.825, FOXG1 P=0.003, NES P=0.796 (* p≤0.05, ** p≤0.01, *** p≤0.001).

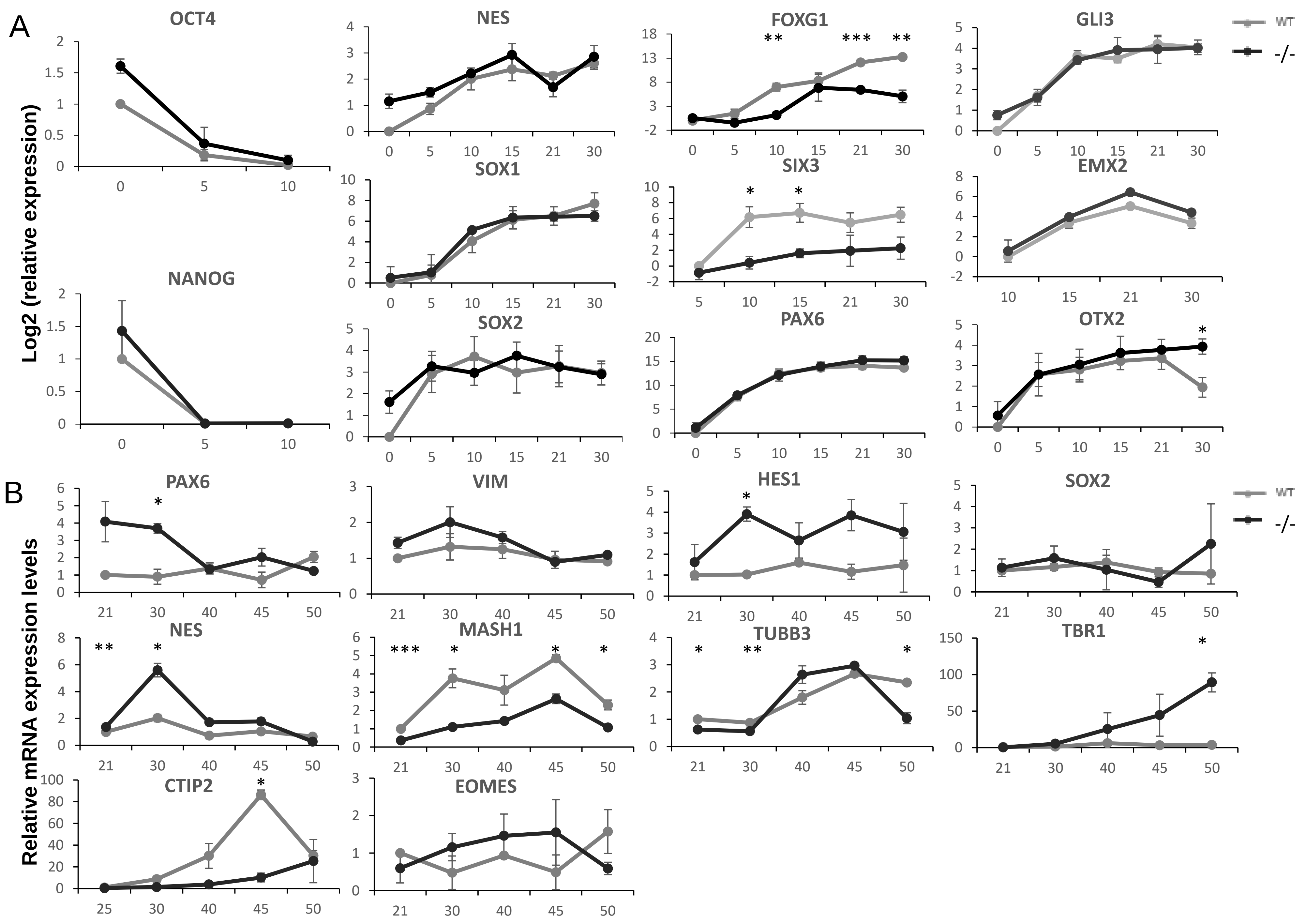

A
B
OCT4
NES
FOXG1
GLI3
SOX1
SIX3
EMX2
NANOG
SOX2
PAX6
OTX2
VIM
HES1
MASH1
TUBB3
TBR1
CTIP2
EOMES
WT
-/-
Log2 (relative expression)
Relative mRNA expression levels

**Fig. S3. Validation of cortical induction by qPCR.** (**A**) qPCR analysis of pluripotency and telencephalic marker genes. RNA samples were harvested every 5 days from day 0 to 30. Levels of mRNA expression were normalized to day 0 mRNA levels. SETBP1-/- levels were normalized against WT levels. Data shown are Log2RQ levels. (**B**) qPCR analysis of neural stem cells and neuronal marker genes. RNA samples were harvested every 5 days from day 21 to 50. Levels of mRNA expression were normalized to day 21. SETBP1-/- levels were normalized against WT levels. Data shown are RQ levels. Data shown as mean ± s.e.m of two independent differentiations analyzed in triplicates. Student's T test was used to compare the expression between the two lines (* $p \leq 0.05$, ** $p \leq 0.01$, *** $p \leq 0.001$).

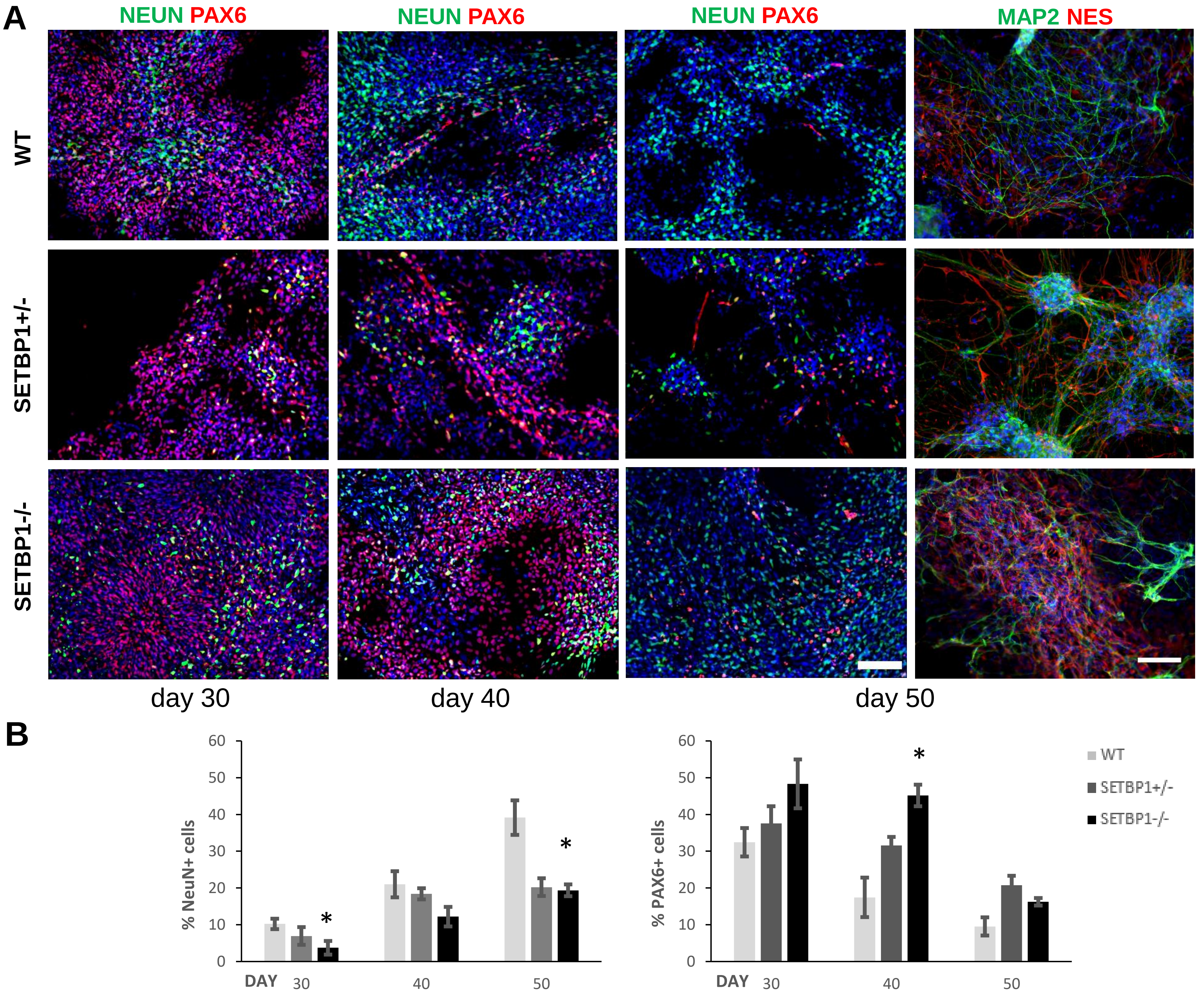
A
NEUN PAX6
NEUN PAX6
NEUN PAX6
MAP2 NES
WT
SETBP1+/-
SETBP1-/-
day 30
day 40
day 50
B
% NeuN+ cells
% PAX6+ cells
DAY 30 40 50
WT
SETBP1+/-
SETBP1-/-
*

**Fig. S4. Expression of NPC and neuronal markers.** (**A**) Cultures were immunostained for PAX6 and NeuN at day 30, 40 and 50, and NES and MAP2 at day 50, Dapi was used to label all nuclei. Scale bar: 100μM. (**B**) Quantitative data for NeuN positive cells (day 30 P=0.036, day 40 P=0.136, day 50 P=0.018) and PAX6 positive cells (day 30 P=0.108, day 40 P=0.032, day 50 P=0.060). Quantitative data presented as mean ± s.e.m for each genotype with a minimum of two independent experiments carried out per line (WT = 5, HET2 = 2, KO1 = 3, KO2 and KO3 = 2). Student's T test was used to compare the expression between the lines (* $p \leq 0.05$, **$p \leq 0.01$, ***$p \leq 0.001$).

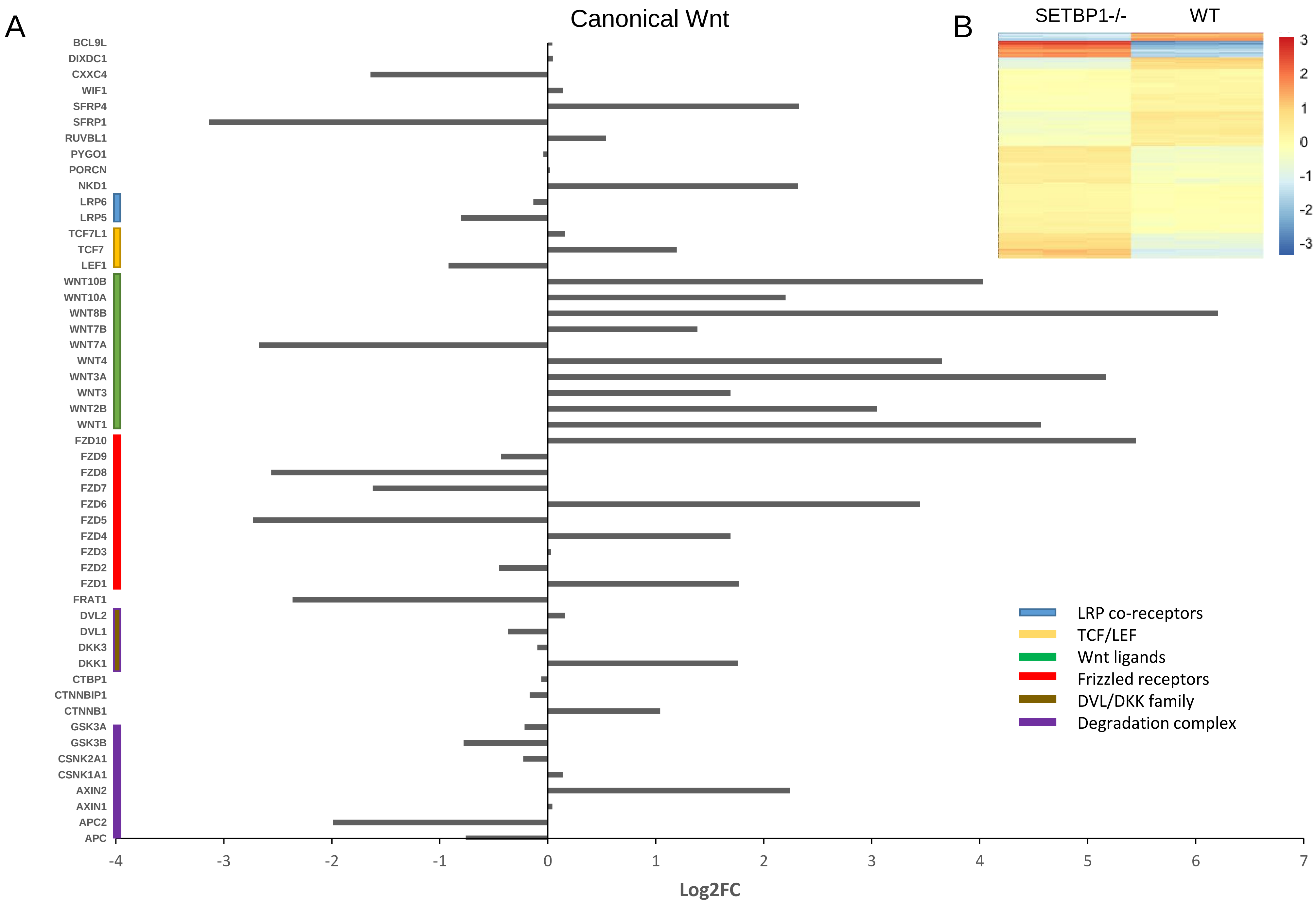
A
Canonical Wnt
B
SETBP1-/-
WT
3
2
1
0
-1
-2
-3
BCL9L
DIXDC1
CXXC4
WIF1
SFRP4
SFRP1
RUVBL1
PYGO1
PORCN
NKD1
LRP6
LRP5
TCF7L1
TCF7
LEF1
WNT10B
WNT10A
WNT8B
WNT7B
WNT7A
WNT4
WNT3A
WNT3
WNT2B
WNT1
FZD10
FZD9
FZD8
FZD7
FZD6
FZD5
FZD4
FZD3
FZD2
FZD1
FRAT1
DVL2
DVL1
DKK3
DKK1
CTBP1
CTNNBIP1
CTNNB1
GSK3A
GSK3B
CSNK2A1
CSNK1A1
AXIN2
AXIN1
APC2
APC
-4
-3
-2
-1
0
1
2
3
4
5
6
7
Log2FC
LRP co-receptors
TCF/LEF
Wnt ligands
Frizzled receptors
DVL/DKK family
Degradation complex

**Fig. S5. Altered expression of genes associated with Wnt pathway.** (**A**) Transcriptomic expression of canonical-Wnt related genes represented as Log2FC. (**B**) Heatmap depicting differentially expressed transcripts for Wnt signalling (GO:0016055). Differentially regulated transcripts used in this analysis were restricted to those with adjusted p-value<0.1, FDR of 10% and a FC>1.5. Benjamini-Hochberg correction was applied for multiple comparisons.

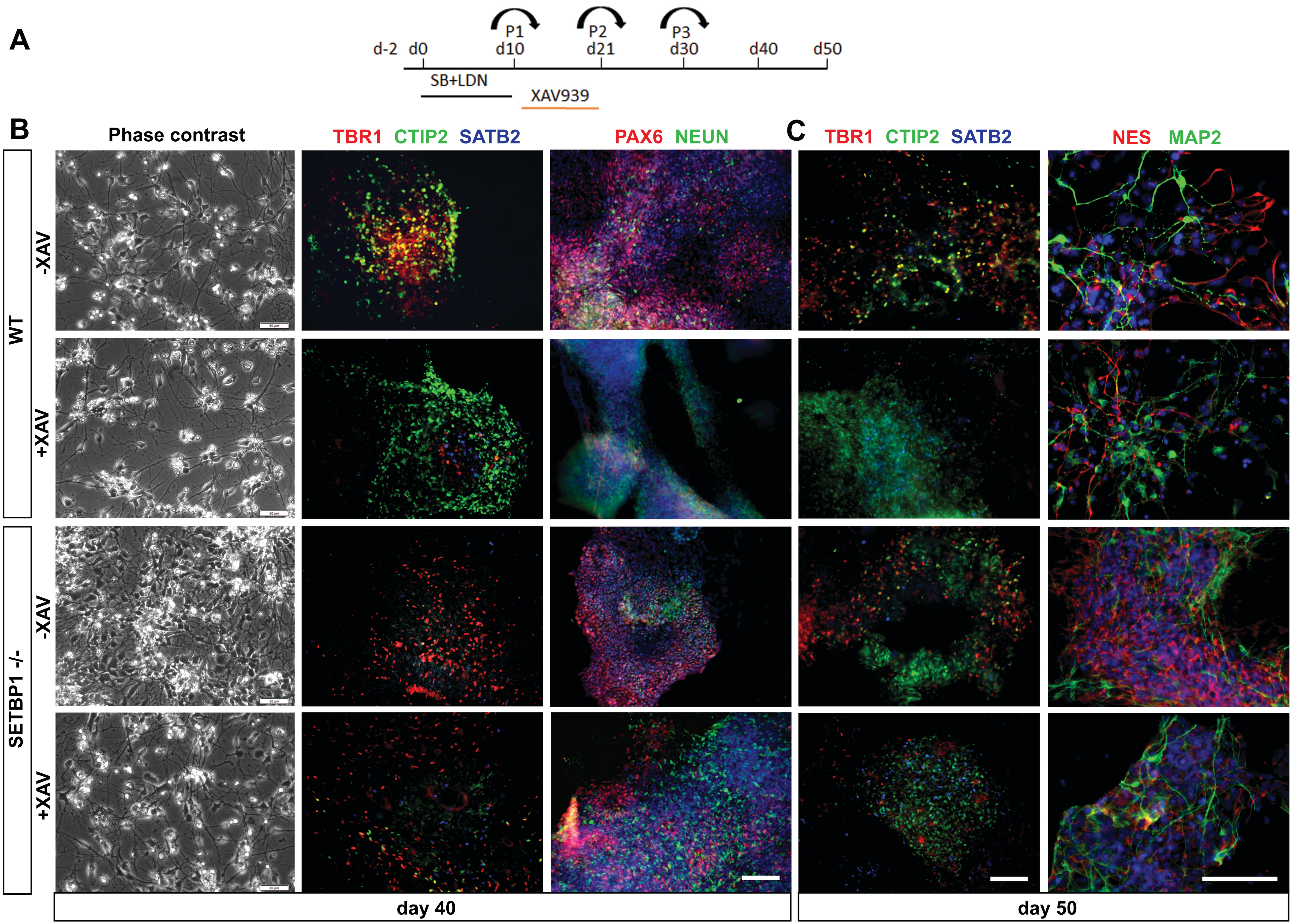
A
P1
P2
P3
d-2
d0
d10
d21
d30
d40
d50
SB+LDN
XAV939
B
Phase contrast
TBR1 CTIP2 SATB2
PAX6 NEUN
C
TBR1 CTIP2 SATB2
NES MAP2
WT
-XAV
+XAV
SETBP1 -/-
-XAV
+XAV
day 40
day 50

**Fig. S6. Phenotype recovery after XAV939 treatment.** (**A**) Experimental scheme indicating window for XAV939 treatment. (**B**-**C**) Phase contrast and fluorescent images of cultures in basal and XAV condition at day 40 and day 50. The blue staining in the 3rd and 5th column are nuclei stained with DAPI. Scale bar: 100uM.

**Table S1. Primers for PCR and qPCR used in the study. qPCR primers were designed to anneal at 60C.**

| Amplicon/ gene | Forward 5'-3' | Reverse 5'-3' | Size (bp) |
|---|---|---|---|
| SETBP1 5'HA | TAAGCGAATTCAAGGCCCAGGGT GAAAGG | CGAATGTCGAC CCACCACCGCTTTAATGGAC | 502 |
| SETBP1 3'HA | GTACAGCGGCCGCGGGCTTTGC TCAGACAC | ACATGGGATCCTTTAGGGACCTGC AGGAC | 551 |
| NHEJ gRNA1 | GCTGAGTAGCGCAGACAAA | CTTTTTCCTTGGCCTGTGTC | 180 |
| NHEJ gRNA2 | CCGGCAAAAGCATCTCATTG | TGGCACCGGGTAATAGTGAT | 240 |
| 5'HA Nested 1 | CAGAATTGCTTCATCAGTCCAG | GCTCGTAGAAGGGGAGGTTG | 1713 |
| 5'HA Nested 2 | CTGGCAGAGAAACTGCAAGC | GGCTTGTACTCGGTCATGGTA | 1157 |
| 3'HA Nested 1 | AGCTGCAAGAACTCTTCCTCAC | GACACAGTTATAAGGTCTGCTGG | 1531 |
| 3'HA Nested 2 | CGATGATCTAGAGCTCGCTGA | CTCCTTCCACTTAGTGCACG | 944 |
| OCT4 | CGACCATCTGCCGCTTTGAG | CCCCCTGTCCCCCATTCCTA | 573 |
| NANOG | AGCCTCTACTCTTCCTACCACC | TCCAAAGCAGCCTCCAAGTC | 278 |
| FOXG1 | TGGCCCATGTCGCCCTTCCT | GCCGACGTGGTGCCGTTGTA | 77 |
| GLI3 | GCTCCACGACCACTGAAAAG | CTGTCCAGGACTTTCATCCTCATTA | 125 |
| OTX2 | TGCCAAAAAGAAGACATCTCCA | AAGCTGGGCTCCAGATAGACAC | 137 |
| SIX3 | CAACTGGTTTAAGAACCGGC | TTACCGAGAGGATGGAGGTG | 218 |
| PAX6 | AGAGAATACCAACTCCATC | GATAATGGGTTCTCTCAAG | 152 |
| EMX1 | ACCGGAGGACAAAGTACAAAC | TAGTCATTGGAGGTGACATCG | 148 |
| EMX2 | GCTTCTAAGGCTGGAACACG | CCAGCTTCTGCCTTTTGAAC | 149 |
| NES | AGCAGGAGAAACAGGGCCTAC | CTCTGGGGTCCTAGGGAATTG | 218 |
| SOX1 | GGAAGGTCATGTCCGAGGCC | ACTTGTCCTTCTTGAGCAGCG | 138 |
| SOX2 | CATGGCAATCAAAATGTCCA | TTTCACGTTTGCAACTGTCC | 102 |
| VIM | CGGGAGAAATTGCAGGAGGA | AAGGTCAAGACGTGCCAGAG | 105 |
| HES1 | GTGTCAACACGACACCGGAT | GGAATGCCGCGAGCTATCTT | 161 |
| EOMES | CTGCCTACCAAAACACCGATATTA | AGCGGGCTTGAGGTAAAGTG | 168 |
| NEUROG1 | GAGACCTGCATCTCCGACCT | TCAGACGCCCGGGAGATATT | 164 |
| MASH1 | GTCCTGTCGCCCACCATCTC | CCCTCCCAACGCCACTGAC | 251 |
| TBR1 | GACTCAGTTCATCGCCGTCA | GCCGGTGTAGATCGTGTCAT | 149 |
| CTIP2 | CTCCGAGCTCAGGAAAGTGTC | TCATCTTTACCTGCAATGTTCTCC | 129 |
| TUBB3 | CATGGACAGTGTCCGCTCAG | CAGGCAGTCGCAGTTTTCAC | 175 |
| GAPDH | ATGACATCAAGAAGGTGGTG | CATACCAGGAAATGAGCTTG | 177 |

**Table S2. Top 20 differentially expressed KEGG pathways in day 15 dataset.**

| Term | Count | % | PValue | Fold Enrichment | Benjamini |
|---|---|---|---|---|---|
| hsa05206:MicroRNAs in cancer | 24 | 2.421796 | 8.17E-05 | 2.449849 | 0.02093 |
| hsa04550:Signaling pathways regulating pluripotency of stem cells | 22 | 2.21998 | 3.14E-04 | 2.353488 | 0.039838 |
| hsa04974:Protein digestion and absorption | 15 | 1.513623 | 4.53E-04 | 2.906977 | 0.038394 |
| hsa04512:ECM-receptor interaction | 16 | 1.614531 | 4.54E-04 | 2.778617 | 0.028981 |
| hsa04151:PI3K-Akt signaling pathway | 38 | 3.834511 | 5.19E-04 | 1.782946 | 0.026536 |
| hsa04390:Hippo signaling pathway | 22 | 2.21998 | 0.001458 | 2.101329 | 0.061059 |
| hsa05205:Proteoglycans in cancer | 25 | 2.522704 | 0.003273 | 1.867611 | 0.114226 |
| hsa04360:Axon guidance | 19 | 1.917255 | 0.005315 | 2.016427 | 0.158476 |
| hsa05410:Hypertrophic cardiomyopathy (HCM) | 12 | 1.210898 | 0.007884 | 2.468694 | 0.203696 |
| hsa04066:HIF-1 signaling pathway | 14 | 1.412714 | 0.010567 | 2.176852 | 0.240539 |
| hsa05200:Pathways in cancer | 39 | 3.935419 | 0.012302 | 1.47737 | 0.252816 |
| hsa05414:Dilated cardiomyopathy | 12 | 1.210898 | 0.01364 | 2.292359 | 0.256521 |
| hsa04510:Focal adhesion | 24 | 2.421796 | 0.015136 | 1.671512 | 0.262032 |
| hsa04310:Wnt signaling pathway | 17 | 1.715439 | 0.018268 | 1.863324 | 0.289006 |
| hsa04261:Adrenergic signaling in cardiomyocytes | 17 | 1.715439 | 0.022539 | 1.818605 | 0.325396 |
| hsa05160:Hepatitis C | 15 | 1.513623 | 0.023104 | 1.910299 | 0.315036 |
| hsa04514:Cell adhesion molecules (CAMs) | 15 | 1.513623 | 0.023104 | 1.910299 | 0.315036 |
| hsa05202:Transcriptional misregulation in cancer | 17 | 1.715439 | 0.025781 | 1.789965 | 0.328288 |
| hsa04068:FoxO signaling pathway | 16 | 1.614531 | 0.026347 | 1.828662 | 0.318997 |
| hsa05217:Basal cell carcinoma | 9 | 0.908174 | 0.026912 | 2.456099 | 0.310563 |

**Table S3. Top 20 differentially expressed KEGG pathways in day 21 dataset.**

| Term | Count | % | PValue | Fold Enrichment | Benjamini |
|---|---|---|---|---|---|
| hsa05217:Basal cell carcinoma | 15 | 1.582278 | 3.26E-06 | 4.439103 | 7.81E-04 |
| hsa04550:Signaling pathways regulating pluripotency of stem cells | 24 | 2.531646 | 4.32E-06 | 2.931217 | 5.18E-04 |
| hsa04310:Wnt signaling pathway | 22 | 2.320675 | 3.72E-05 | 2.730287 | 0.002975 |
| hsa04390:Hippo signaling pathway | 24 | 2.531646 | 3.74E-05 | 2.582751 | 0.00224 |
| hsa05322:Systemic lupus erythematosus | 15 | 1.582278 | 6.20E-05 | 3.497475 | 0.00297 |
| hsa04080:Neuroactive ligand-receptor interaction | 26 | 2.742616 | 8.02E-05 | 2.353595 | 0.003204 |
| hsa04916:Melanogenesis | 17 | 1.793249 | 1.54E-04 | 2.939451 | 0.00526 |
| hsa04020:Calcium signaling pathway | 22 | 2.320675 | 3.71E-04 | 2.334866 | 0.011066 |
| hsa04512:ECM-receptor interaction | 15 | 1.582278 | 4.61E-04 | 2.921941 | 0.012216 |
| hsa04640:Hematopoietic cell lineage | 11 | 1.160338 | 4.72E-04 | 3.761728 | 0.011277 |
| hsa04911:Insulin secretion | 14 | 1.476793 | 7.97E-04 | 2.911411 | 0.017256 |
| hsa05200:Pathways in cancer | 39 | 4.113924 | 0.001576787 | 1.66713 | 0.031068 |
| hsa04924:Renin secretion | 10 | 1.054852 | 0.003686359 | 3.14059 | 0.065909 |
| hsa00512:Mucin type O-Glycan biosynthesis | 7 | 0.738397 | 0.004372125 | 4.308889 | 0.072363 |
| hsa04360:Axon guidance | 17 | 1.793249 | 0.006408089 | 2.092889 | 0.097746 |
| hsa04610:Complement and coagulation cascades | 8 | 0.843882 | 0.008558683 | 3.327327 | 0.120967 |
| hsa04261:Adrenergic signaling in cardiomyocytes | 17 | 1.793249 | 0.01229605 | 1.952322 | 0.160264 |
| hsa04974:Protein digestion and absorption | 11 | 1.160338 | 0.012612234 | 2.453301 | 0.155687 |
| hsa04022:cGMP-PKG signaling pathway | 18 | 1.898734 | 0.014450806 | 1.871622 | 0.167955 |
| hsa04151:PI3K-Akt signaling pathway | 29 | 3.059072 | 0.018060931 | 1.554975 | 0.196447 |

**Table S4. Top 20 differentially expressed KEGG pathways in day 34 dataset.**

| Term | Count | % | PValue | Fold Enrichment | Benjamini |
|---|---|---|---|---|---|
| hsa04510:Focal adhesion | 43 | 4.35663627 | 1.43E-10 | 2.94444588 | 3.51E-08 |
| hsa05205:Proteoglycans in cancer | 39 | 3.95136778 | 3.30E-09 | 2.84957481 | 4.06E-07 |
| hsa04512:ECM-receptor interaction | 24 | 2.43161094 | 9.53E-09 | 3.97331172 | 7.81E-07 |
| hsa04390:Hippo signaling pathway | 30 | 3.03951368 | 7.28E-07 | 2.74380792 | 4.48E-05 |
| hsa04141:Protein processing in endoplasmic reticulum | 32 | 3.24214792 | 7.43E-07 | 2.63221489 | 3.65E-05 |
| hsa04974:Protein digestion and absorption | 19 | 1.92502533 | 1.17E-06 | 3.7651142 | 4.81E-05 |
| hsa05217:Basal cell carcinoma | 16 | 1.62107396 | 3.30E-06 | 4.1031585 | 1.16E-04 |
| hsa04310:Wnt signaling pathway | 24 | 2.43161094 | 4.78E-05 | 2.53138408 | 0.00146736 |
| hsa04151:PI3K-Akt signaling pathway | 42 | 4.25531915 | 5.40E-05 | 1.90732759 | 0.00147431 |
| hsa04916:Melanogenesis | 18 | 1.82370821 | 2.09E-04 | 2.73742697 | 0.00513827 |
| hsa05146:Amoebiasis | 17 | 1.72239108 | 2.39E-04 | 2.81442913 | 0.00533209 |
| hsa04550:Signaling pathways regulating pluripotency of stem cells | 22 | 2.2289767 | 5.93E-04 | 2.2479218 | 0.0120836 |
| hsa05206:MicroRNAs in cancer | 22 | 2.2289767 | 0.00133263 | 2.1156911 | 0.02491859 |
| hsa04670:Leukocyte transendothelial migration | 17 | 1.72239108 | 0.00175984 | 2.3653181 | 0.03047617 |
| hsa05166:HTLV-I infection | 30 | 3.03951368 | 0.00250845 | 1.77540512 | 0.04035341 |
| hsa05200:Pathways in cancer | 43 | 4.35663627 | 0.00323646 | 1.55786472 | 0.04861961 |
| hsa00512:Mucin type O-Glycan biosynthesis | 7 | 0.70921986 | 0.00965075 | 3.66206897 | 0.13092887 |
| hsa04810:Regulation of actin cytoskeleton | 24 | 2.43161094 | 0.01280928 | 1.69671149 | 0.16154222 |
| hsa00910:Nitrogen metabolism | 5 | 0.50658561 | 0.01369796 | 5.03031451 | 0.16354159 |
| hsa05412:Arrhythmogenic right ventricular cardiomyopathy | 12 | 1.21580547 | 0.01437938 | 2.274577 | 0.16318337 |
